# Chandelier cell anatomy and function reveal a variably distributed but common signal

**DOI:** 10.1101/2020.03.31.018952

**Authors:** Casey M. Schneider-Mizell, Agnes L. Bodor, Forrest Collman, Derrick Brittain, Adam A. Bleckert, Sven Dorkenwald, Nicholas L. Turner, Thomas Macrina, Kisuk Lee, Ran Lu, Jingpeng Wu, Jun Zhuang, Anirban Nandi, Brian Hu, JoAnn Buchanan, Marc M. Takeno, Russel Torres, Gayathri Mahalingam, Daniel J. Bumbarger, Yang Li, Tom Chartrand, Nico Kemnitz, William M. Silversmith, Dodam Ih, Jonathan Zung, Aleksandar Zlateski, Ignacio Tartavull, Sergiy Popovych, William Wong, Manuel Castro, Chris S. Jordan, Emmanouil Froudarakis, Lynne Becker, Shelby Suckow, Jacob Reimer, Andreas S. Tolias, Costas Anastassiou, H. Sebastian Seung, R. Clay Reid, Nuno Maçarico da Costa

## Abstract

The activity and connectivity of inhibitory cells has a profound impact on the operation of neuronal networks. While the average connectivity of many inhibitory cell types has been characterized, we still lack an understanding of how individual interneurons distribute their synapses onto their targets and how heterogeneous the inhibition is onto different individual excitatory neurons. Here, we use large-scale volumetric electron microscopy (EM) and functional imaging to address this question for chandelier cells in layer 2/3 of mouse visual cortex. Using dense morphological reconstructions from EM, we mapped the complete chandelier input onto 153 pyramidal neurons. We find that the number of input synapses is highly variable across the population, but the variability is correlated with structural features of the target neuron: soma depth, soma size, and the number of perisomatic synapses received. Functionally, we found that chandelier cell activity *in vivo* was highly correlated and tracks pupil diameter, a proxy for arousal state. We propose that chandelier cells provide a global signal whose strength is individually adjusted for each target neuron. This approach, combining comprehensive structural analysis with functional recordings of identified cell types, will be a powerful tool to uncover the wiring rules across the diversity of cortical cell types.

## Introduction

Inhibition powerfully shapes the activity of neuronal circuits. In the mammalian neocortex, the simplest role of inhibition is to restrain the network from “runaway” excitation, but the diversity of inhibitory cell types, each with distinctive projection patterns and physiology, suggest a far richer role in cortical computation (Ascoli et al., 2008; Fino et al., 2013; Freund and Buzsáki, 1996; Jiang et al., 2015; Kepecs and Fishell, 2014; Kubota, 2014). Understanding the contribution of an individual cell type to circuit function requires knowledge of both the underlying rules governing synaptic connectivity and the conditions under which it is functionally active. For example, the finding that some vasoactive intestinal polypeptide (VIP)-positive interneurons preferentially inhibit somatostatin (SST)-expressing interneurons (Pfeffer et al., 2013) and are active during locomotion (Fu et al., 2014) strongly suggested an important role in the state-dependent modulation of cortical processing (McGinley et al., 2015; Reimer et al., 2014; Vinck et al., 2015). Similar efforts have uncovered circuit roles of other cell types, including parvalbumin (PV)-expressing basket cells (Packer and Yuste, 2011; Wilson et al., 2012) and SST-expressing Martinotti cells (Silberberg and Markram, 2007; Wang et al., 2004; Wilson et al., 2012).

The Chandelier cell (ChC) has unique properties that, at first glance, should cast it to be one of the better-understood inhibitory cell types. Sometimes referred to as axo-axonal cells, ChCs are a class of GABAergic interneurons characterized by a number of vertical axonal “candles” or “cartridges” (Jones, 1975; Peters et al., 1982; Szentágothai and Arbib, 1974) that synapse exclusively with the axon initial segment (AIS) of excitatory pyramidal neurons (PyCs) (DeFelipe et al., 1985; Fairén and Valverde, 1980; Somogyi, 1977; Somogyi et al., 1982). This pattern of connectivity suggests a unique role for ChCs, as the AIS is a specialized compartment approximately 10–40 *µ*m from the axon hillock whose unique ion channel distribution makes it the principal site of action potential generation (Kole and Stuart, 2012; Kole et al., 2007; Palmer, 2006). Each ChC makes synapses onto 30-50% of PyCs within a 200 *µ*m wide axonal domain (Wang et al., 2019), and thus a low density of overlapping of cells can potentially innervate all PyCs in L2/3 (Inan et al., 2013) and thus have the potential for tremendous impact on cortical activity.

Despite their highly specific connectivity, the ways in which ChCs affect neuronal circuits remain surprisingly enigmatic (Inan and Anderson, 2014; Woodruff et al., 2010). One reason is the variability in the strength of ChC targeting. PyC populations in different brain regions and layers receive diverse numbers of AIS-targeting boutons (DeFelipe et al., 1985; Veres et al., 2014; Wang and Sun, 2012). Consistent with this observation, activation of the ChC population *in vivo* has diverse effects on nearby PyCs, from strong inhibition to no response (Lu et al., 2017), despite nearly all PyCs likely receiving ChC input (Inan et al., 2013). The logic underlying this variability, and thus the heterogeneity of ChC influence on the downstream targets, is largely unknown, although categorical differences in long-range projection targets are one potential factor (Lu et al., 2017). A second reason is functional. What conditions drive ChC activity remains unknown. ChCs receive synaptic input from local PyCs (Jiang et al., 2015; Lu et al., 2017), Martinotti cells (Jiang et al., 2015), cholinergic projections from basal midbrain (Lu et al., 2017), and gap junctions from other ChCs (Woodruff et al., 2011), but how these and other unknown inputs conspire to drive activity in a behaving animal is unclear. Adding to the complexity, while GABA is typically inhibitory in mature animals, the biophysics of the AIS can result in ChCs depolarizing their targets (Szabadics et al., 2006; Woodruff et al., 2009, 2011), although ChCs are functionally inhibitory under the more typical condition of coincident excitatory input (Woodruff et al., 2011).

To improve our understanding of both the circuit organization and function of ChCs, we took a multipronged approach. We used large-scale serial-section electron microscopy (EM) (Bock et al., 2011; Kasthuri et al., 2015; Lee et al., 2016) to map AIS input across L2/3 PyCs in a volume of mouse primary visual cortex (Supplemental Video 1). By using automated dense segmentation and synapse detection, we obtained a reconstruction of ChC axons and PyCs in a volume of approximately 3.6 10^6^ *µ*m^3^. The resolution and completeness afforded by this approach allowed us to infer underlying principles governing not only of the presence, but also the structural weight of ChC connectivity. To address ChC function, we used a genetic line that selectively expresses in ChCs to record neuronal activity in awake behaving mice. We used these results to inform biophysical modeling of ChC inputs on individual pyramidal neurons to explore what makes such inhibition unique.

## Results

### A densely segmented EM volume of layer 2/3 primary visual cortex

A single large EM volume of cortical tissue reveals the cellular and subcellular components, from neuronal morphology to synaptic connectivity and organelles. The ability to map the fine ultrastructure of all cells across hundreds of microns can be used to study numerous questions across neuroscience and cell biology. Thus, with broad utility in mind, we prepared a volumetric EM image volume of (L2/3) of mouse primary visual cortex (Figure 1A) spanning approximately 250 *µ*m × 140 *µ*m × 90 *µ*m with 40 nm thick sections imaged at 3.58 × 3.58 nm/pixel with transmission EM (Figure 1B; see methods for details).

**Figure 1.**
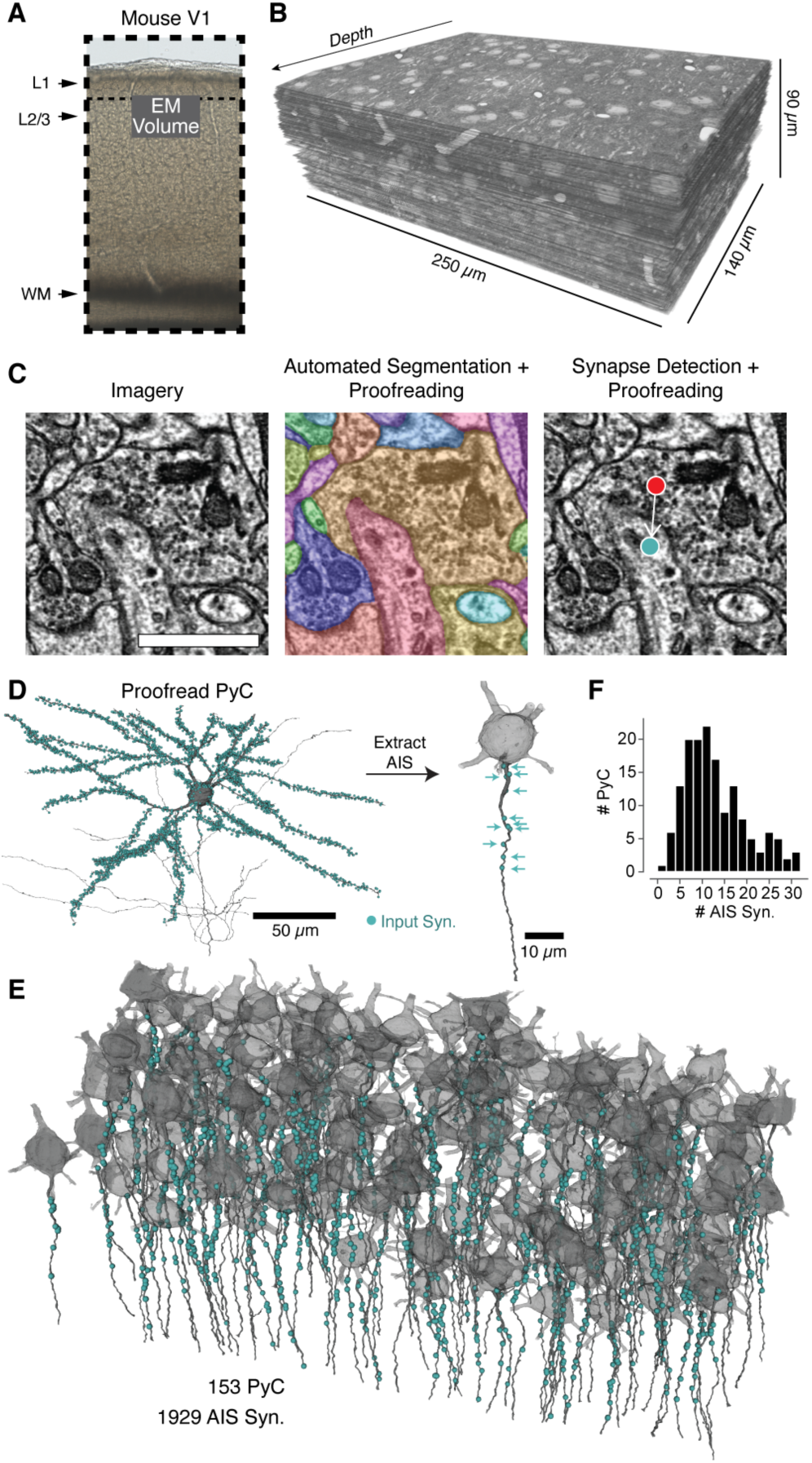
A map of AIS input from EM. A) A block of tissue was selected from L2/3 of mouse V1 and processed in an EM pipeline. B) Serial 40 nm sections were imaged and computationally aligned. C) Image annotation pipeline. Left, images were taken at 3.58 × 3.58 nm/pixel resolution. Ultrastructure and membrane staining is clear. Scale bar is 500 nm. Center, the neuropil was densely segmented and targeted proofreading was done to correct PyCs and other objects of interest. Right, automated synapse detection identified pre- and post-synaptic locations for synapses and was followed by targeted proofreading for false positives. D) For each PyC with sufficient AIS in the volume, we started with the full morphology and all synaptic inputs (cyan dots). We then computationally extracted the AIS (dark gray) and its synaptic inputs (cyan arrows). E) Soma, AIS and AIS synaptic inputs for all PyC analyzed. The volume is rotated so that the average AIS direction is downward. Note that dendrites and higher order axon branches are omitted for clarity. F) Histogram of synapses per AIS.

In order to address neuronal connectivity, we needed to be able to follow a multitude of individual axons throughout the volume. Such circuit reconstruction at scale is intractable without intensive computational processing (Berning et al., 2015; Dorkenwald et al., 2017; Jain et al., 2010; Januszewski et al., 2018). We used a series of novel machine learning based methods to perform high-quality image alignment, automated segmentation, and synapse detection for the volume (Dorkenwald et al., 2019). Nonetheless, proofreading is still necessary for precise measurements of anatomy and connectivity. The initial segmentation identified small supervoxels that were agglomerated into cells, and we built a novel cloud-based proofreading system to edit the agglomerations to perform targeted error correction. As a basis to begin analysis, the 458 cell bodies in the volume were manually classified as excitatory, inhibitory, or glia based on morphology and ultrastructural features and all 364 PyCs were proofread to correct segmentation errors (Dorkenwald et al., 2019).

### A complete map of synaptic input to the axon initial segments of an excitatory network

While ChCs are the only cell type to specifically target the AIS, other cell types can also form AIS synapses (Gonchar et al., 2002; Kisvárday et al., 1985; Somogyi, 1977). To narrow down our search for ChC inputs, we first mapped all synaptic input onto the AIS of all excitatory cells that had a complete AIS in the volume (N=153). Since we cannot robustly identify the molecular markers that define the AIS, we opted instead for a purely structural definition: from the axon hillock (whether it emerged from the soma or a proximal dendrite) to the most proximal of the first branch point, beginning of myelination, or the volume exit (if at least 40 *µ*m of AIS was within the volume, a distance found to contain almost all AIS synapses, see Figure 2I). For each PyC, we manually marked these two points as the top and bottom of the AIS. We used these points to computationally specify the AIS and its synaptic input for each labeled PyC (Figure 1D,E, Supplemental Figure 1A) and proofread for false-positive synapse detections. This resulted in a total of 1929 AIS synapses across the 153 PyCs, for a mean of 12.6 synapses per AIS. However, the mean hides a remarkable diversity of total input, from 1-32 synapses per AIS (Figure 1F), suggesting large differences in AIS inhibition.

**Figure 2.**
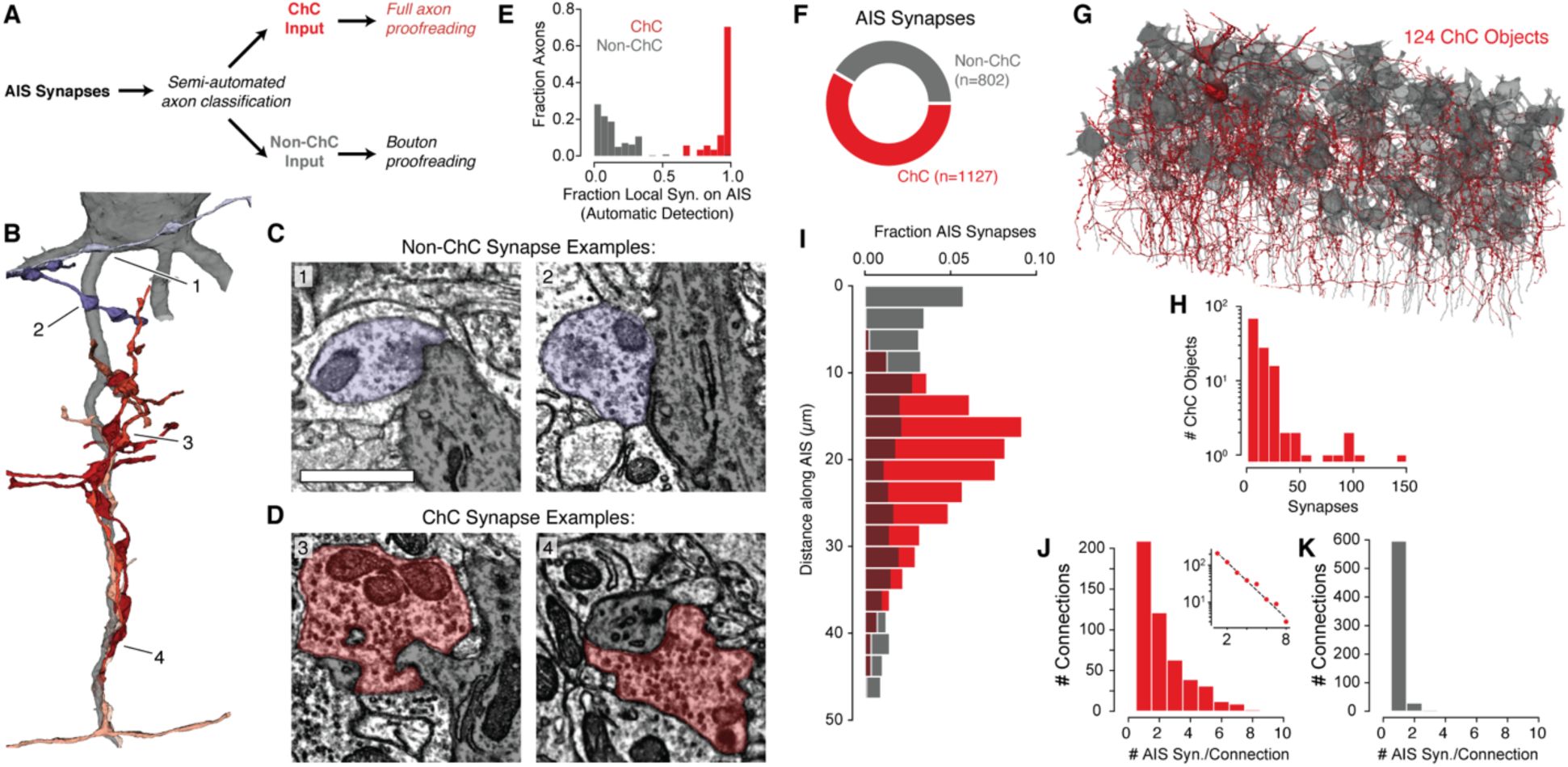
Characterization of AIS inputs. A) Classification and proofreading workflow. Across AIS synaptic inputs, morphology and connectivity were used to distinguish ChC from non-ChC axons. ChC axons were extensively proofread to get as-complete-as-possible arbors. For non-ChC inputs, all AIS-targeting boutons were proofread to ensure there were no falsely merged ChC axons. B) 3d morphology of a typical AIS and the axons that innervate it. Non-ChC axons are in shades of purple, ChC axons are in shades of red. Different colors indicate distinct axons. Note that ChCs form chains of boutons and cluster together on the AIS. Numbers correspond to the panels in C and D. C and D) In single sections, both non-ChC (C, purple) and ChC (D, red) boutons are unambiguous synapses, but can look similar from local imagery alone. E) Histogram of fraction of automatically detected AIS synapses onto analyzed PyCs for each presynaptic axon before synapse proofreading. Note that almost all non-AIS synapses from ChCs were false positives (see text). For clarity, axons with three or more synapses are shown. F) Breakdown of AIS synapse count by axon class. G) Morphology of all distinct ChC objects (red) and PyCs (gray). Note two ChC somata in the volume, with the rest being orphan axon fragments of varying sizes. H) Distribution of output synapses per distinct ChC object. I) Distribution of the distance from the soma of AIS synapses by axon class. J) Distribution for ChCs of the number of AIS synapses per connection (i.e. synapse count from an axon object to an AIS). Inset shows the same data on a semilog-y scale with a geometric fit (black dashed line). K) Distribution for non-ChCs of the number of AIS synapses per connection.

### PyC AIS input is a mix of Chandelier and non-Chandelier synapses

We next sought to identify which AIS inputs came from ChCs and which did not (Figure 2A). Using the dense segmentation, we examined the morphology (Figure 2B) and ultrastructure (Figure 2C,D) of every axon presynaptic to any AIS synapse. All but two axons that targeted AISs were orphan fragments (*i.e.* they did not connect to a cell body within the volume). However, using the EM reconstructions, we distinguished ChC axons from non-ChC using the degree of AIS-selectivity, propensity to form multi-bouton cartridges, and fine scale anatomy (see Methods). ChC axons were extensively proofread to extend their morphology to be as complete as possible within the volume. Non-ChC axons were proofread only to the extent necessary to classify them.

We validated our classification by looking at how AIS-targeting axons synapsed onto PyC compartments. Since every PyC input could be labeled AIS or non-AIS, we considered all automatically detected synapses from AIS targeting axons onto the PyCs (n=15,376). As expected, ChC axons consistently and overwhelmingly formed AIS synapses, while non-ChC axons preferred other compartments (Figure 2E). Many non-ChC axons had morphology and connectivity consistent with soma-targeting basket cells. Our automated synapse detection also found ChC synapses onto other compartments, but most were found to be false positives (42/49, i.e. not a synapse), while for the remainder the ultrastructure was ambiguous.

In total, we found 1127 AIS synapses from 124 ChC axon fragments and 802 AIS synapses from 612 non-ChC axon fragments (Figure 2F, G). Individual ChC axons ranged in size, with between 1–150 presynaptic sites in the volume (Figure 2H). We found that ChC synapses were preferentially located on the AIS in a region between 10–40 *µ*m from the axon hillock, while non-ChC synapses were widely distributed but more common nearest to the soma (Figure 2I). It is thus likely that much of the non-ChC inputs were located within the transition between the somatic and AIS compartments, while ChCs have more specialized targeting (Tai et al., 2019).

Given the low density of ChC cells in cortex, we were surprised to find two Chandelier soma with both axon and dendrites in the volume (Figure 2G). Connections from other neurons within the volume onto both ChC cells were observed, including synaptic input from both local PyCs and a putative L2/3 Martinotti cell, as well as putative dendro-dendritic gap junctions between the ChC cells (Supplementary Figure 2).

### Distribution of AIS inputs by cell type

An AIS can have inputs from many different ChCs, with each connection typically thought to be a multi-bouton “cartridge”. For clarity, we use “synapse” to refer to a single anatomical synapse and “connection” to indicate the collection of synapses between a given presynaptic and postsynaptic neuron, comprising one or more synapses. Here, the ability to discriminate each presynaptic axon gives the unique opportunity to consider the properties of how a complete map of synapses is organized by connection. While it is possible for a single ChC to use multiple axonal branches to target a given AIS, whole-cell reconstructions suggest this is rare (Blazquez-Llorca et al., 2015; Gouwens et al., 2019). We therefore assume that each ChC fragment targeting the same AIS comes from a distinct cell.

We first asked about the properties of the connections between a presynaptic axon and an AIS. The synapse count for individual ChC connections ranged from one to nine and was well-fit by a geometric distribution (exponent: 0.44, Figure 2J). Notably, this included both standard multi-synapse connections that characterize ChC axon morphology, as well surprisingly numerous single-synapse connections. Counting only cartridges would have undercounted total ChC synapses by 20%. In contrast, non-ChC inputs were found throughout the AIS, but most commonly near the soma. Many non-ChC axons also targeted the same cell’s soma, and we speculate that this initial AIS region is often treated by basket cell axons as a continuation of the somatic compartment. Non-ChC connections comprised only a single synapse in 94% of examples (no non-ChC connection had more than 3 synapses). While they could potentially contribute to AIS inhibition, individual non-ChC connections are on average much weaker than ChC connections.

### Chandelier cell input to pyramidal cells is highly variable and correlates with other forms of inhibition

We next asked how ChC input was distributed across the population of L2/3 PyCs. Because different PyCs had different amounts of axonal arbor in the volume, we restricted our analysis to a consistent initial region that would both cover the typical molecularly defined AIS and include as many synapses as possible. Based on the distribution of ChC synapses (Figure 2I), we settled on using the first 37 *µ*m of structural AIS for all cells, which contained 97% of ChC synapses (Figure 3A).

**Figure 3.**
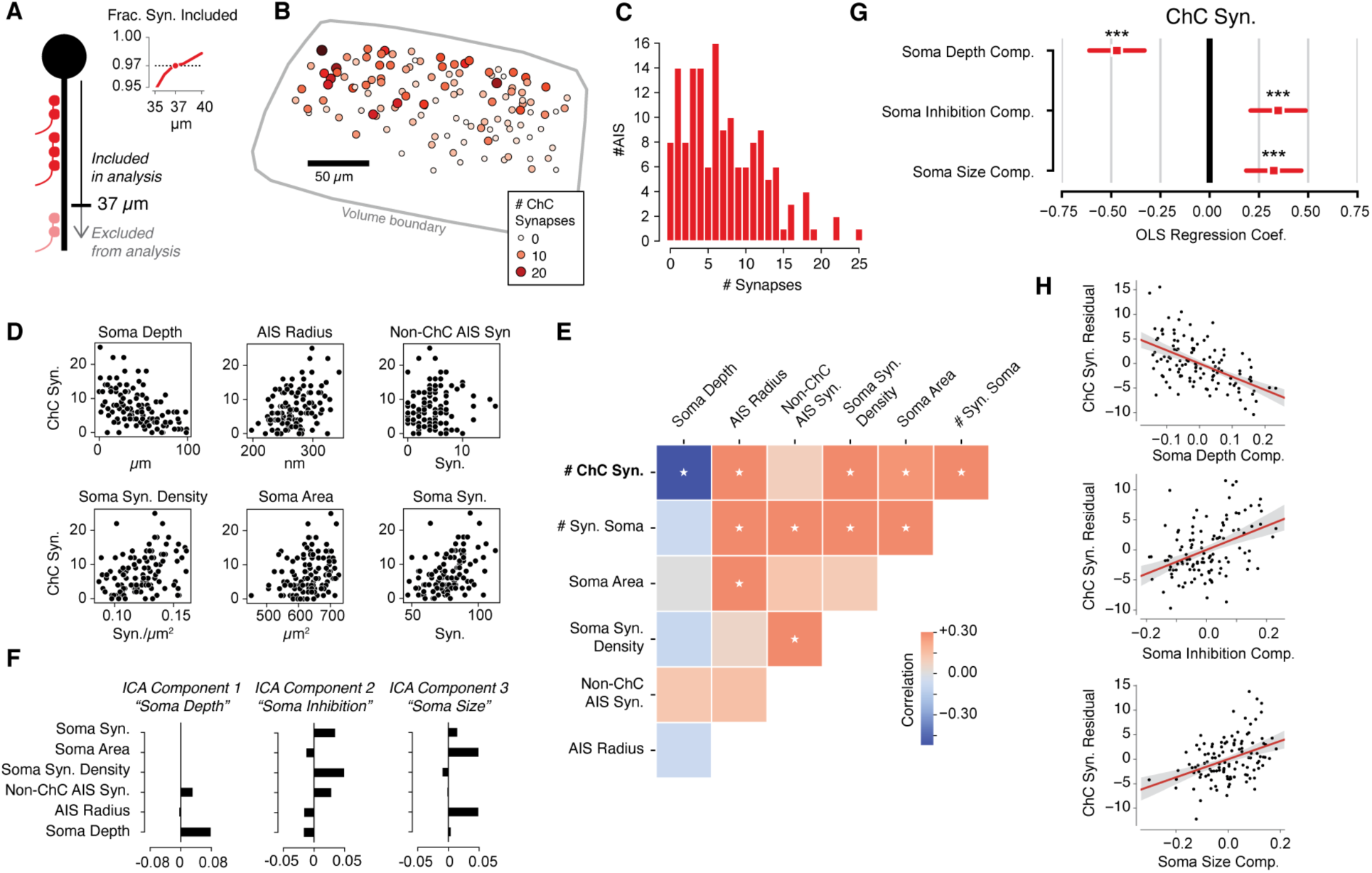
Structural factors affecting ChC input on PyCs. A) Schematic of AIS analysis. To compare all PyCs evenly, we focused only on those synapses within the first 37 microns from the axon hillock. Inset: Fraction of all observed synapses included in analysis by threshold. B) Soma location and ChC synapse count for each analyzed PyC, with volume boundary (gray outline). Pia is up. C) Histogram of ChC synapse count per PyC. D) ChC synapse count versus somatic and AIS structural properties: soma depth (measured from the shallowest cell analyzed), average AIS radius, non-ChC AIS synapse count, Soma synapse density, Soma surface area, and soma synapse count. E) Pearson correlation between ChC synapse count and structural properties, as well as structural properties with one another. White dots indicate p<0.05 to be different from zero after Holm-Sidak multiple test correction. F) ICA components for PyC structural properties. Note that each component has most of its contribution from an interpretable combination of properties. G) Regression coefficients of ChC synapse count (z-scored) on ICA components. Bars indicate 95% confidence interval before multiple correction, stars indicate significance after Holm-Sidak multiple test correction. ***: p<0.001. H) Scatterplots and linear fit for each structural property versus the residual of ChC synapse count after fitting the other two properties. Shaded region is the 95% confidence interval from bootstrapping.

ChC inputs were found on 95% of PyCs (144/151), but there was striking variability in the total ChC input (Mean: 7.4 ± 5.4 synapses) with individual PyCs receiving between 0–25 ChC synapses (Figure 3B, C). Effectively, some L2/3 PyCs escape ChC inhibition entirely while others have the potential to be strongly inhibited. This was not the case for other sources of perisomatic inhibition. Using similar methods as we used to find AIS synapses, we computationally identified the synapses onto the PyCs’ soma. All PyCs had numerous somatic inputs (47–113 synapses, Figure 3D).

To explore the logic of this heterogeneity, we asked if the amount of ChC input that a PyC receives could be associated with other structural properties of the cell. We focused on properties relating to the soma and the AIS, as those could be completely observed for most PyCs (n=114, see Methods and Supplemental Figure 1). Specifically, for each PyC we measured its depth within L2/3, mean AIS radius, number of non-ChC AIS synapses, number of synaptic inputs onto the soma, soma surface area, and soma synapse density. Strikingly, we found significant correlations between the number of ChC inputs and each property other than non-ChC AIS synapses (Figure 3D,E). However, we also found that various size and synaptic input properties of each PyC were significantly correlated among themselves (Figure 3E). To disentangle these correlations, we performed Independent Components Analysis (ICA), yielding three interpretable mixtures of variables representing the soma depth, somatic inhibition (i.e. soma synapses, soma synapse density, and non-ChC AIS synapses) and soma size (i.e. soma area and AIS radius) (Figure 3F). Importantly, as a mathematical consequence of ICA, these PyC structural components are fully uncorrelated with one another (Supplemental Figure 3).

To gain insight into ChC input variability, we performed multivariate ordinary least-squares regression (OLS) on the number of ChC synapses against the three PyC structural components. Despite the simplicity of the approach, linear combinations of perisomatic structural properties explained 45% of the variance in ChC input, with significant effects for all three components (Figure 3G, H). We report coefficients from z-scored variables. First, deeper PyCs received less ChC inhibition (Coefficient: −0.47), which is consistent with the observation that some ChC axons have denser axonal arbors in upper L2/3 (Wang et al. 2019) and our observation of a higher number of ChC synapses in the upper part of our volume than the lower (Supplemental Figure 4). Second, perisomatic inhibition was positively correlated with ChC synapses (Coefficient: 0.33), meaning that PyCs with more ChC input also received more input from other, non-ChC cells at their soma. Third, larger cells received more ChC synapses (Coefficient: 0.34). Taken together, the magnitude of ChC input is extremely diverse and influenced by multiple aspects of the target cells.

**Figure 4.**
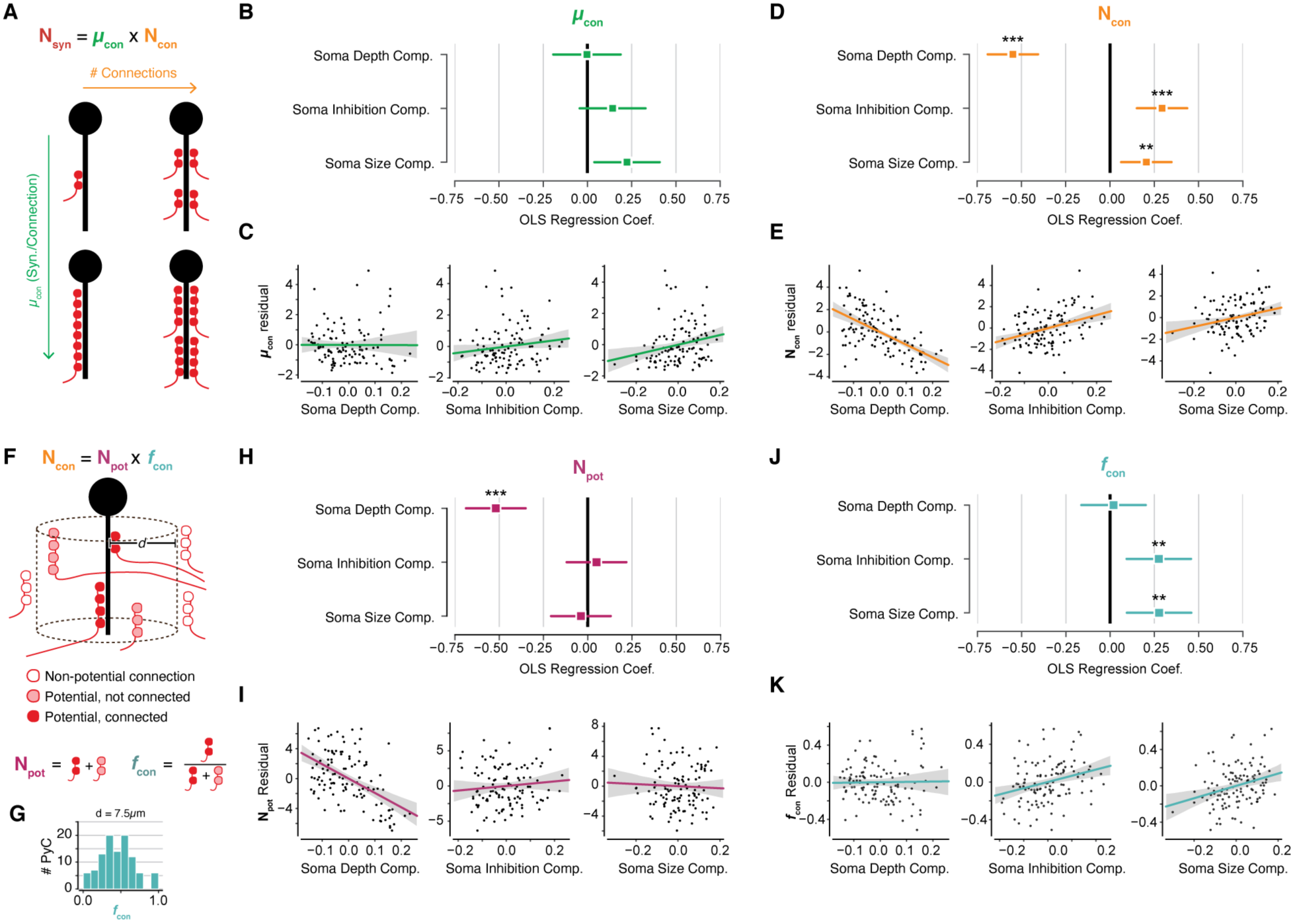
Breaking ChC input into constituent aspects. A) The total ChC synapse count for an AIS is, trivially, the product of the number of connections and the mean number of synapses per connection. Either or both aspects could correlate with PyC structural properties to modulate ChC input. B) Regression coefficients of the average number of synapses per connection against PyC structural components. No values are significant after multiple testing correction. C) Scatterplots and linear fit for each structural property versus the residual of mean synapses per connection after fitting the other two properties. Note that trends match those of the overall synapse count. D) OLS analysis for the number of ChC connections. E) Residuals as in C, for the number of ChC connections. F) To measure the geometric effect on number of connections, we measure potential connections as those within a distance d of an AIS and the connectivity fraction as the number of potential connections actually connected. G) Histogram of connectivity fraction for d=7.5*µ*m, the distance threshold used in H–K. See Supplemental Figure 5 for higher and lower distance thresholds. H,I) OLS analysis for potential connections. Note that potential connections are only associated with depth. I) Residuals as in C, for potential connections. J) OLS analysis for connectivity fraction. Note that connectivity fraction is only associated with the soma inhibition and soma size components, but not depth. K) Residuals as in C, for connectivity fraction. In the OLS coefficient plots, the lines indicate the 95% confidence interval. Stars indicate significance after Holm-Sidak multiple test correction. **: p<0.01, ***: P<0.001. For residual plots, the shaded region is the 95% confidence interval after bootstrapping.

### Cell-specific factors influence the number of Chandelier synapses on the axon initial segment

We next sought to better understand the nature of the wide variability in ChC input. We first considered the different biologically distinct scenarios that can yield the same overall number of ChC inputs onto a PyC. The same number of ChC synapses onto a single PyC can be produced by different combinations of number of connections and number of synapses per connection. (Figure 4A). Since each ChC axon was individually segmented in the EM data, we were able to measure both components for every PyC. While variability in total synapses could be due to either factor individually, we found that both ranged widely (1-9 connections and 1-7 synapses per connection). However, a similar OLS approach as above found no significant relationship between synapses per connection and PyC structural properties (Figure 4B, C), while the number of connections had a similar relationship to all PyC structural properties as total ChC synapses (Figure 4D,E). This suggests that the number of connections and the number of synapses per connection are regulated by different processes.

Variability in the number of connections could be due to geometry alone, for example if there were more ChC axons per AIS in some parts of the volume than others. To test this, we used a simple model to account for spatial effects based on (Stepanyants et al., 2002). We define “potential connections” as those ChC-PyC pairs where a ChC axon and AIS come within a given distance, and “connectivity fraction” as the fraction of potential connections that are associated with synapses (Figure 4F). The number of connections can then be decomposed into the number of potential connections times the connectivity fraction. If input variability were determined only by local abundance of ChC axons, then we would expect potential connections but not connectivity fraction to be related to PyC structural properties.

Using the geometry of every ChC axon fragment that innervates any of the AISs, we computed both potential connections and connectivity fraction for each PyC. To account for unobserved ChC axons past the edge of the volume, we omitted those AISs too close to volume boundaries (see Methods). Since these measurements depend on the distance threshold and we lack a detailed understanding of ChC axon guidance or target finding, we looked for distance thresholds that had a wide span of connectivity fractions between 0–1. We show results for a distance threshold of 7.5 *µ*m (Figure 4G), although the same results hold for thresholds between 5-10 *µ*m (Supplemental Figure 5). Using the same OLS approach as above, we found that while both potential connections and connectivity fraction showed significant relationships with PyC structural properties, different properties mattered for each. The number of potential connections decreased with depth but had no significant relationship with soma size or soma inhibition (Figure 4H,I). In contrast, connectivity fraction was not significantly affected by soma depth, but was positively modulated by soma size and soma inhibition components (Figure 4J,K). Together, these results indicate that while soma location modulates the local abundance of ChC axons around a given AIS, there are also additional cell-specific factors that affect the recruitment of ChC axons to an AIS.

**Figure 5.**
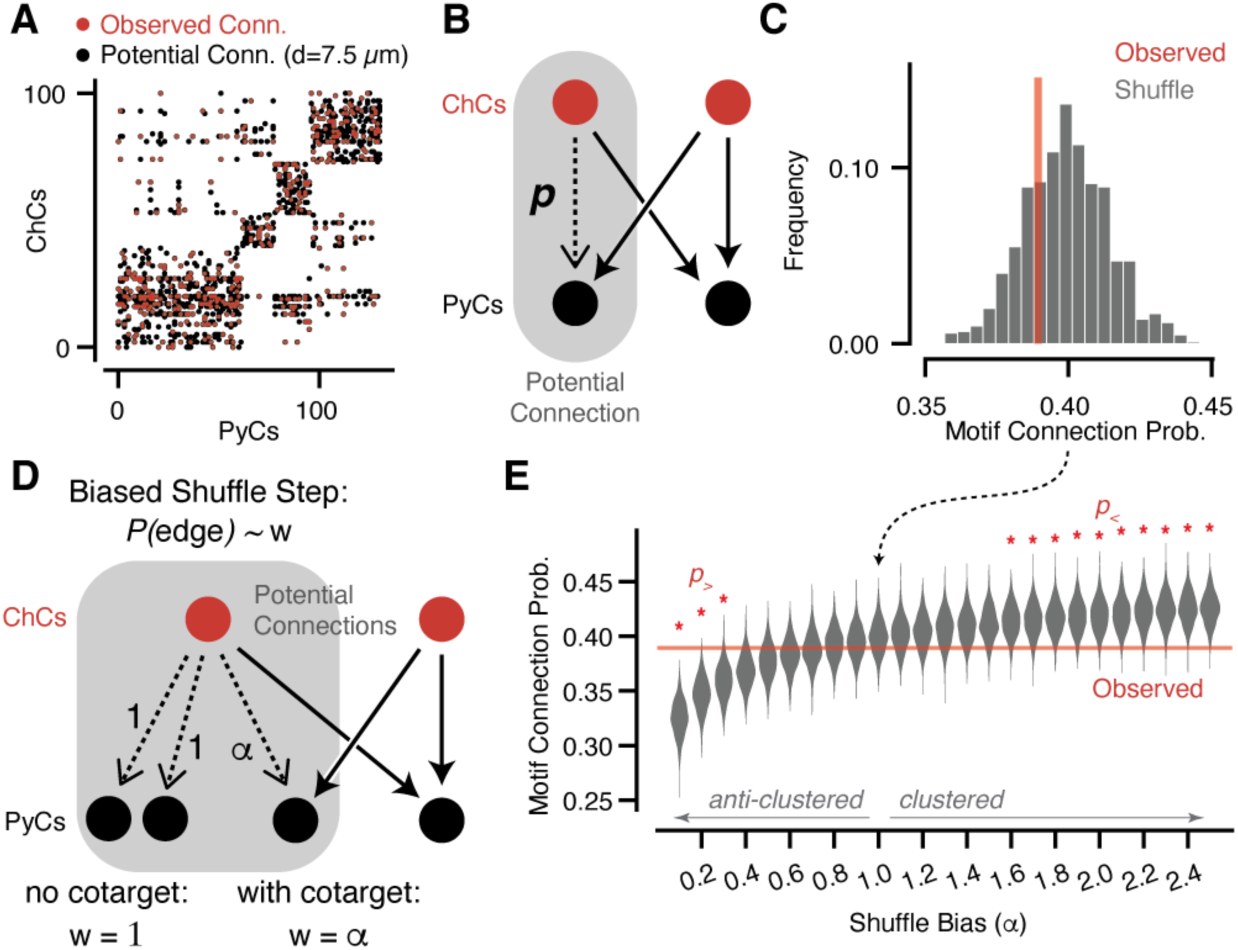
No evidence of clustering of ChC targets based on geometrically-constrained motif analysis. A) Connectivity matrix from ChC axons onto PyC targets. Each black dot represents one potential connection for a distance threshold of 7.5 *µ*m (see Supplemental Figure 6 for alternative thresholds)) and each red dot represents one actual connection with any number of synapses. Elements are clustered by the potential connectivity matrix, suggesting that most of the structure of the network comes from geometry alone. B) Cartoon of a potential bifan motif. In a bifan motif, two ChCs would target the same pair of PyCs. In a potential bifan motif, three of those connections are present and the fourth is a potential connection. We consider the probability p that this potential edge is connected. C) Observed connectivity probability for the potential bifan motif in our dataset (red line) compared to shuffled networks that preserve PyC in-degree and potential connectivity. We see no evidence of excess clustering of ChC targets beyond geometry. D) Cartoon of a biased bifan shuffle, where potential connections that complete a bifan motif are given a different weight in the shuffle probability. E) Observed connectivity (red line) probability versus shuffled networks with different bias weights based on the shuffle step shown in D (gray violin plots, N=1000 per bias weight value). Red stars indicate bias weights where the observed value is above (p>) or below (p>) 95% of the shuffled distribution. Even with the geometric constraints, we can rule out strong clustering or anti-clustering within the data.

### Chandelier connections show no evidence of target selectivity beyond spatial proximity

So far, we have implicitly considered all ChC axons to be part of a uniform population. However, groups of ChC axon fragments could show clustered connectivity, showing a tendency to co-innervate the same set of PyC AISs. If so, it would suggest that ChCs are forming specific inhibitory subnetworks and are not, in fact, a uniform population. While there are numerous clustering measures and community detection algorithms for abstract graphs (Fortunato, 2010), we have the additional need to account for the spatial relationships between neurons, otherwise proximity alone can introduce clustering that is irrelevant to our question (Figure 5A). To account for both clustering and geometry, we measured the connectivity fraction of potential edges only for those that could complete a bifan motif, i.e. one where two ChCs target the same two PyCs (Figure 5B). If ChC input were clustered, this probability should be higher than that observed in networks where ChC connections are randomly shuffled amongst all the potentially connected ChC axons while holding fixed the number of ChC connections for each PyC. The observed probability we measured was well within the range of values obtained by shuffling (Figure 5C), and thus there is no evidence that ChC axons are coordinated beyond geometrical factors that constrain potential connectivity.

To understand how strong a propensity for co-targeting we could detect given our dataset, we performed a power analysis that added a motif-dependent bias to the shuffle. During the shuffle, potential edges that could complete a bifan motif were selected with a weight *α*, while those that did not were selected with weight 1. Thus, *α* > 1 produces networks with increased clustering, *α* < 1 biases against clustering, and *α =* 1 is random (as above). Our simulations found that the observed data would deviate significantly from randomized data for *α* < 0.4 or *α* > 1.5, and thus the data is only consistent with, at most, a modest targeting bias. Larger reconstructions, which would more completely fill in the potential connectivity graph, would likely constrain these bounds further.

### *In vivo* function of Chandelier Cells shows collective activity during periods of arousal

The synapse count between ChCs and PyCs only determines their potential to influence their targets. The functional effect of inhibition depends also on the typical pattern of ChC activity during behavior. If all ChC cells were active at the same time, total synapse count would offer a good approximation of net functional strength (Veres et al., 2014) while if subpopulations of ChCs were activated at different times, the exact pattern of ChC co-activation would be important.

To label chandelier cells *in vivo*, we used a mouse line in which recombinase CRE was coexpressed with Vipr2, a genetic marker expressed specifically in ChCs (Vipr2-IRES2-Cre, Daigle et al., 2018). Full transgenic strategies do not label ChCs in mouse primary visual cortex effectively, likely due to off-target expression during development (Tasic et al., 2018). To circumvent this shortcoming, we injected an AAV viral vector containing a CRE-dependent calcium indicator gene GCaMP6f (Chen et al., 2013) into V1 of adult Vipr2-IRES2-Cre mice. Histological examination showed this strategy specifically labelled neurons at the layer1/layer 2 border with arbors characteristic of ChC morphology (Figure 6A), indicating a faithful labeling of ChCs.

**Figure 6.**
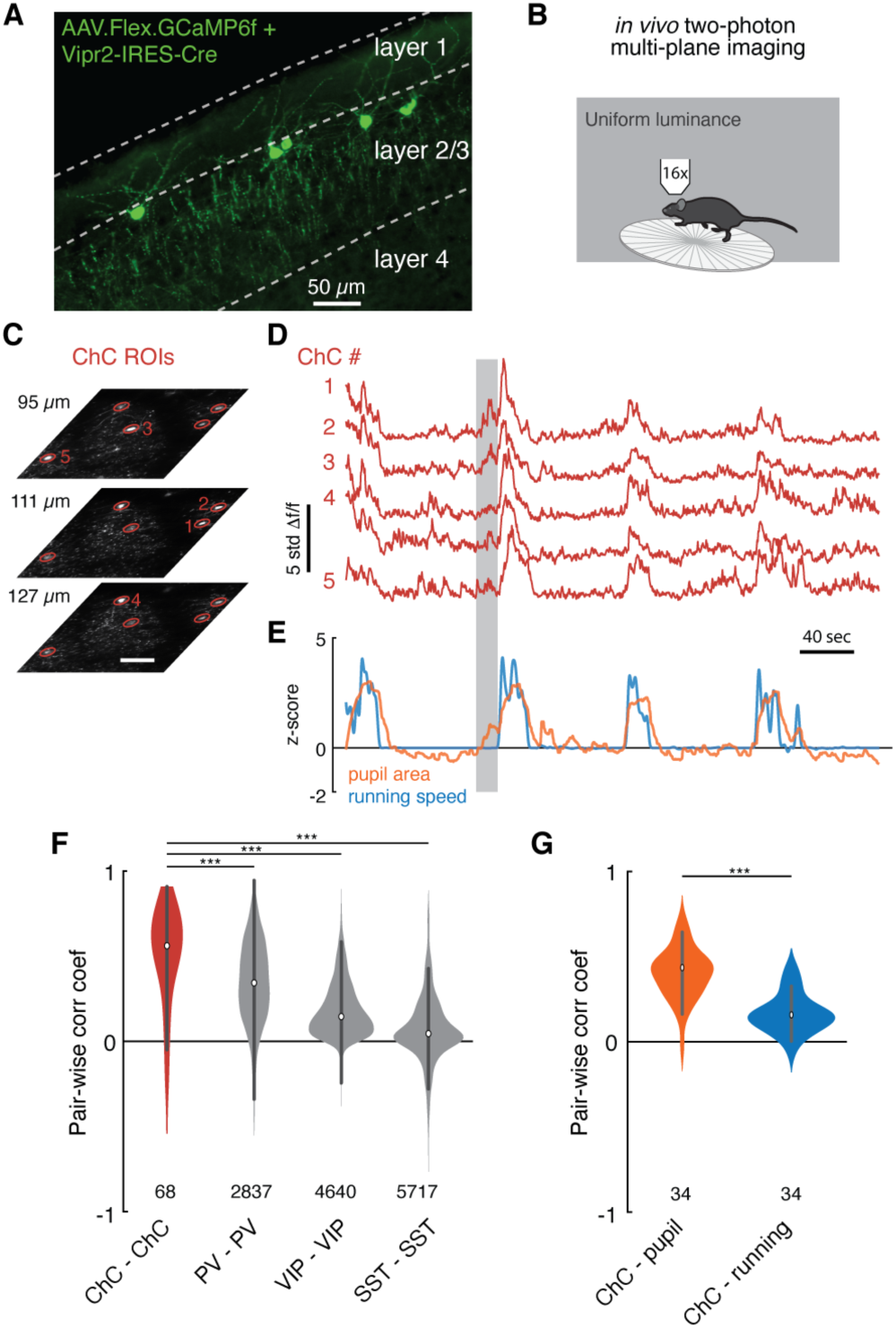
Functional imaging of ChCs reveals a correlated response to arousal state. A) Maximum projection image in upper layers of V1 showing ChC-specific GCaMP6f expression in Vipr2-IRES2-Cre mice injected with AAV-Flex-GCaMP6f. Note cell bodies at the L1/L2 border, characteristic cartridges, and L1 dendrites. B) Cartoon of experimental design. Mice expressing GCaMP6f in V1 ChCs were placed on a treadmill and imaged with multiplane two photon microscopy while subject to a uniform luminance visual stimulus. C) An experiment with simultaneous three plane imaging (depth shown to left) with five distinct ChC ROIs, each measured from its best plane. Scale bar is 50 *µ*m. D) GCaMP6f responses (z-scored from baseline) for the same five ROIs as in C. Note extremely correlated large events. E) Simultaneous behavioral measurements for the same experiment as in D. Increased pupil size, a measure of arousal state, and running bouts correspond to periods of high ChC activity. Note that there are periods where ChC activity and pupil area increase in the absence of running (e.g. the period noted by the gray box). F) Pairwise Pearson correlation for spontaneous activity traces for different interneuron classes. ChC cells measured from two imaging sessions each for three mice show high spontaneous correlation compared to other classes of interneurons, as computed from comparable observations in public Allen Institute Brain Observatory data. Number of distinct pairs is shown. G) Pairwise correlations between ChC activity traces and behavioral traces shows that ChC are significantly more correlated with pupil area than with running. Number of distinct pairs is shown. Statistical significance for cell-cell correlations was computed with the Mann-Whitney U test, while for matched cell-behavior correlations significance was computed with the Wilcoxon signed rank test. ***: p<0.001.

Using this strategy, we measured the *in vivo* calcium activity of ChCs by two photon imaging (in mice not imaged for EM). To simultaneously acquire imagery from sparsely distributed cells (Figure 6A), we used a multi-plane imaging system (Liu et al., 2018) that allowed near-simultaneous monitoring across a range of cortical depth (Figure 6C). The imaging procedure followed a standardized awake behaving paradigm (de Vries et al., 2020), during which head-fixed mice were presented with a screen with uniform luminance and allowed to engage in spontaneous behavior. In all imaging sessions, we observed striking seconds-long bouts where all ChCs were active at the same time. This suggested that not only can ChCs be spontaneously active, but they are often active (or inactive) as a uniform population (Figure 6D). To quantify this, we calculated the average cell-cell correlation between ChC cells during spontaneous behavior (Figure 6E, correlation coefficient, ChC-ChC: 0.49±0.30, 68 pairs). The spontaneous correlation among ChCs was notable even compared to other common interneuron cell types (correlation coefficient, PV-PV: 0.34±0.24, 4640 pairs; VIP-VIP: 0.18±0.16, 5717 pairs; SST-SST: 0.07±0.20, 2837 pairs), as estimated from Allen Brain Observatory data (de Vries et al., 2020) (Figure 6E).

When comparing ChC activities with animals’ behavior states, we noticed that coordinated ChC activities always occurred during periods of dilated pupils and locomotion, indicative of an active arousal state (McGinley et al., 2015; Reimer et al., 2014; Vinck et al., 2015). There were some pupil dilation events in the absence of locomotion that showed concurrent ChC activation (Figure 6E, gray bar). Indeed, ChC cell activities were more strongly correlated with pupil diameter than with locomotion speed (Figure 6G, correlation coefficient, ChC-pupil vs. ChC-locomotion: 0.40±0.14 vs. 0.16±0.11, 34 pairs, T=3.0, p=4.8×10^−7^, Wilcoxon signed-rank test). From these observations, we conclude that during periods of high arousal, L2/3 PyCs receive input from all or nearly all of their ChC synapses.

### Biophysical simulations of ChC inhibition on pyramidal neurons

Given that the activity of ChCs is highly correlated and their synapses are tightly clustered together on the AIS, we wondered how the innervation pattern of multiple ChCs on an AIS affects the activity of PyCs. To address these questions, we used computational single-neuron models, instantiated excitatory and inhibitory input along their detailed morphology, and asked how synaptic properties affect their input-output transformation. We generated biophysically detailed models of V1 pyramidal neurons utilizing an optimization procedure based on whole-cell patch-clamp experiments and associated reconstructed cellular morphology (see Methods). Importantly, the single-cell models developed contain active Na-, K- and Ca-dependent conductances along their entire morphology, including the AIS. This, in turn, results in these models reproducing a number of physiological properties of real neurons such as action potentials initiated at the AIS that are back-propagated to the soma and the dendritic arbor (Figure 7A–C). We used two models from layer 2/3 neurons and two models from layer 4 neurons in order to test if the postsynaptic cell type affected any of our conclusions.

**Figure 7.**
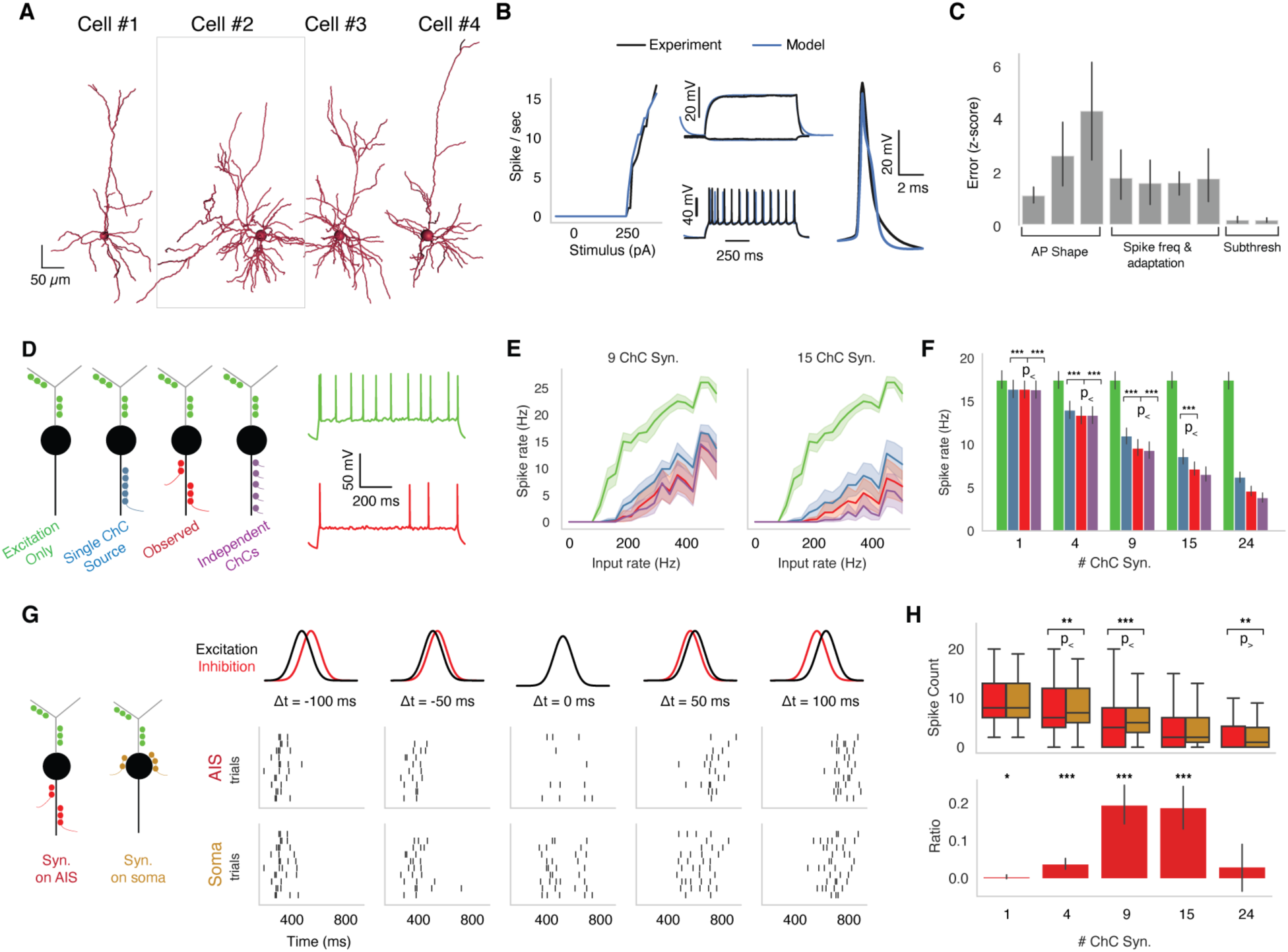
Biophysical modeling of ChC inhibition. **A**, Reconstructed morphologies for four PyCs from mouse V1. **B**, Biophysically-detailed, all-active models were fit for each cell constrained by its electrophysiological features and reconstructed morphology. Panels show model neuronal response and observed data for one cell (#2 in A). Left, F/I curve; middle-top, subthreshold voltage responses; middle-bottom, suprathrehold voltage response; right, mean spike shape. **C**, Average z-scored training errors for the four models for physiological parameters. **D**, Left, schematic for different configurations of ChC synapses. Right, example simulated functional traces for excitation only (green) and observed inhibition (red) in response to a step function stimulus. **E**, Mean F/I curves for each synapse configuration for 9 total ChC synapses and 15 total ChC synapses for one cell (#2). **F**, Average PyC firing rate at fixed excitation across the four synapse configurations. **G**, (Left) We compared the effect of the same number of synapses at the soma and the AIS in a biologically-inspired configuration as a function of excitatory/inhibitory timing. (Right) Raster plot for a representative cell for each case at different temporal offsets between excitatory and inhibitory barrages. **H**, (Top) The spike count for inhibition at soma and at the AIS, (Bottom) The differential effect in inhibition as computed by the difference between Soma and AIS spike counts divided by the sum of the spike counts. The ratio ranges between −1 and 1, with 1 representing inhibition at the axon being 100% more effective than at the soma and 0 meaning no differential effect.

To assess the impact of the temporal coordination of ChC inhibition along the AIS, we considered four scenarios: one with excitation only and three with different temporal patterns of inhibition (Figure 7D). First, in the synchronous inhibition case, all ChC synapses were activated by the same realization of a Poisson process resulting in perfect inhibitory input synchrony. Second, in the asynchronous inhibition case, each ChC synapse was activated by an independent realization of the Poisson process, resulting in co-active ChC input with uncorrelated spike times. Third, in the biologically-inspired inhibition case, clusters of ChC synapses activated synchronously, but each cluster was activated by a different realization of a Poisson process, simulating the activity of multiple presynaptic ChC (each with one or more synapses) converging onto the same AIS. As expected, in the presence of excitatory synaptic input onto PyC dendrites, ChC inhibition reduced spiking (Figure 7E). Asynchronous ChC input was most effective at suppressing PyC activity, while the synchronous ChC input was least effective. The biologically inspired scenario presented an intermediate case between the most and least efficient case of inhibition. These trends remain robust across neurons and when altering the number of ChC synapses along the AIS across PyC models. These results suggest that the observed innervation of the AIS by multiple ChCs is stronger than if all the synapses were provided by a single ChC.

How does the location of ChC synapses affect the integration properties of pyramidal neurons? Intuitively, the fact that ChC synapses are located along the AIS, the location where action potentials are initiated, puts them in a position to heavily influence the input-output relationship of pyramidal neurons, however previous simulations have shown that this effect is mild (Douglas and Martin, 1990). As previous simulations were with morphologies from a different species, we performed simulations over multiple model neurons of mouse V1 neurons described above. To assess the impact of the spatial constellation of ChC inhibition, we chose the biologically inspired connectivity pattern and compared how inhibition alters integration properties depending on whether it is placed along the AIS or at the soma. We simulated Gaussian barrages of dendritic excitation and either somatic or AIS inhibition with different temporal offsets, from inhibition leading excitation by 100 ms to inhibition trailing excitation by 100 ms (Figure 7G). We observe that temporal disparity between the excitatory and inhibitory barrage leads to overall higher spike output compared to when the two barrages coincide, signifying that inhibition along the AIS dampens excitatory drive (in agreement with Figure 7F). Moreover, the location of ChC synapses affected pyramidal output. While for low and high numbers of inhibitory synapses, the location of synapses did not change pyramidal output, for intermediate amounts of inhibition the AIS was on average more effective in suppressing pyramidal spiking (by approximately 25%) (Figure 7H). These results indicate that when the incoming excitation and inhibition barrages are co-active, the axo-axonic inhibition modulated ChC are more effective than the basket-cell-like somatic projections. One of the strengths of our approach is that we used models of multiple neurons, and while the results were significant and align well with the functional recordings, we also find substantial variability from model to model. Given this variability and that the effect of chandelier inhibition on the spiking of PyCs depends on the level of concurrent excitation (Douglas and Martin, 1990), we therefore wonder if there were other reasons for the specific targeting of the AIS besides preventing the activation of sodium conductances.

### Preferential targeting of ChC synapses to cisternal organelles on the AIS

Unlike more typical somatic and dendritic inhibitory synapses, ChC synapses interact with the unique biophysics of the AIS (reviewed in Leterrier, 2018), giving them the potential to have qualitatively different effects than other inhibitory synapses. While AIS-specific chloride transporter and its effect on the GABA reversal potential has been carefully studied with respect to ChC input (Szabadics et al., 2006; Woodruff et al., 2009, 2011), another interesting but less well-studied factor is calcium. The presence of calcium in the AIS can affect the action potential generation and timing (Bender and Trussell, 2009). Moreover, T-type voltage gated Ca^2+^ channels are present at the AIS (Bender and Trussell, 2009) and are de-inactivated by hyperpolarization. The AIS hosts unique calcium-storing structures called cisternal organelles (CO), stacked endoplasmic reticulum specializations associated with a complex assortment of molecular components, including ryanodine receptors and Kv2.1 ion channels (King et al., 2014). The co-localization of GABAergic synapses and COs has been suggested from qualitative observations (Benedeczky et al., 1994; King et al., 2014), however this could occur by chance, since COs and ChC synapses necessarily occupy a limited space.

The detailed 3D reconstructions and images available from EM allowed us to investigate the relationship between these COs and circuit structure more rigorously. We annotated stacked cisternae (Figure 8A), the ultrastructural hallmark of COs (Peters et al., 1968), on ten PyCs chosen to span the observed variation in ChC synapse number. For each PyC, we manually generated a point cloud covering the extent of each CO and computationally mapped these points to the surface of the AIS. Combining this data with the synapse detection, we computed the depth and angular orientation of both COs and ChC synapses for each AIS (Figure 8B, C). COs were typically found only within the initial 30 *µ*m of the AIS (Figure 8D, E), and appeared to frequently co-localize with ChC synapses in multi-synapse clusters (Figure 8D), although there were some COs without synapses and some synapses without COs. To test if the observed proximity could occur by chance, we measured the distances between each ChC and its closest CO point and performed two types of shuffle test. First, we used depth and orientation measurements to simulate the same configuration of CO positions on other AISs and compared mismatched CO/synapse configurations to those observed (Figure 8E). We found that matched configurations had significantly smaller distances between COs and synapses than shuffled configurations. To single out the effect of angular orientation without changing depth, for each AIS we computationally rotated the orientation of its CO points and again computed synapse-to-CO distances (Figure 8F). Zero degrees of rotation had the minimal median distance, and rotations greater than 90° were significantly different than this baseline state. Taken together, this demonstrates that ChC synapses are preferentially located at COs.

**Figure 8.**
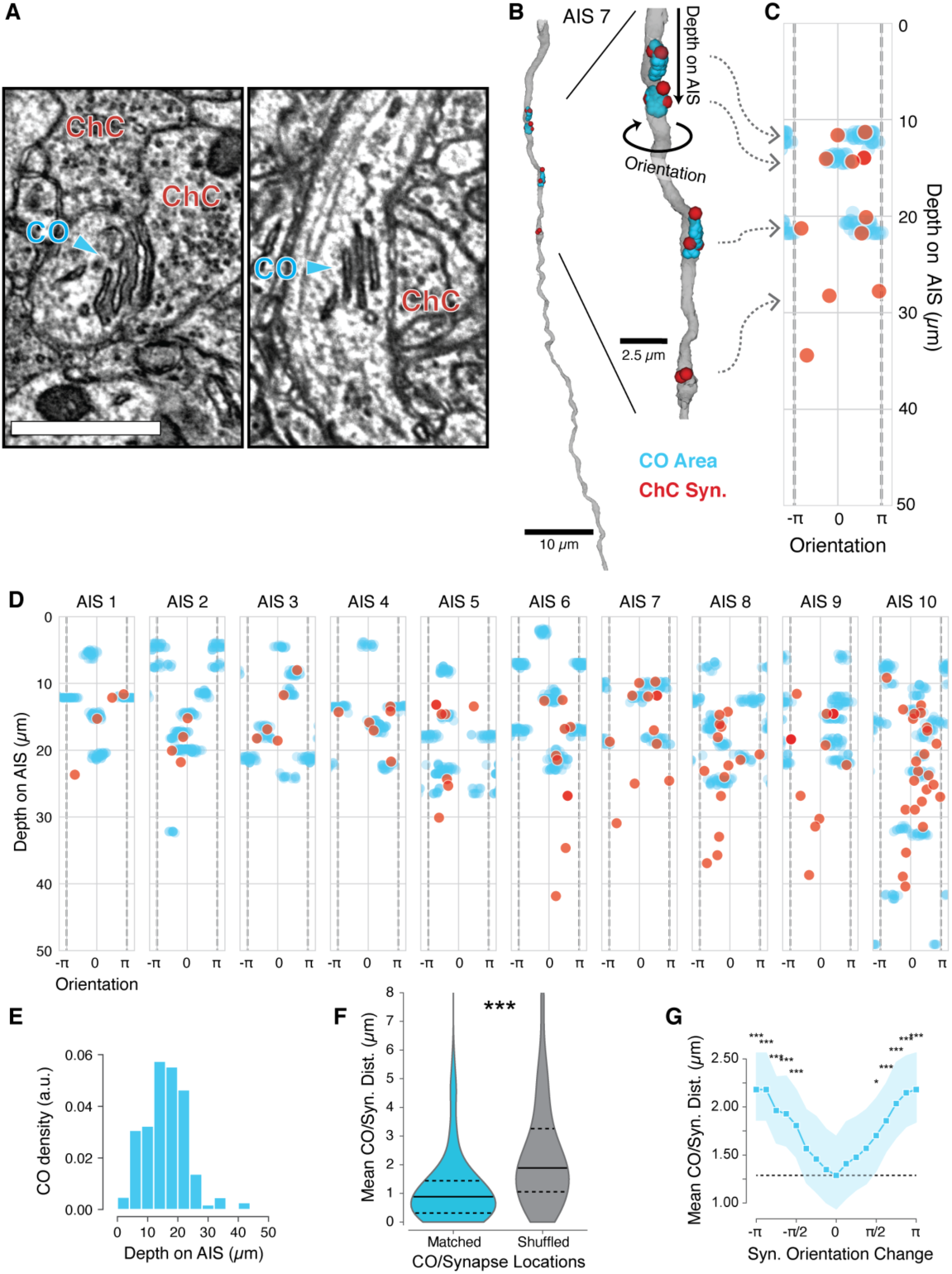
ChCs preferentially target cisternal organelles on the AIS. A) Electron micrograph showing the stacked ER of a cisternal organelle (CO) in an AIS, adjacent to two ChCs. Scale bar is 1 *µ*m. B) Left, 3d view of an AIS (gray) with COs (cyan dots) and ChC synapses (red dots) shown. Right, higher magnification of the same AIS from the region indicated. Depth is measured as the distance from the axon hillock and orientation as the angle around the AIS, with 0 set to the same arbitrary direction for all AISs. C) Representation of the AIS in B in depth and orientation. The COs are shown as cyan point clouds, and the center of ChC synapses are shown as red dots. The plots wrap in orientation at π/-π (gray dashed lines). D) COs and ChC synapses for all ten AISs with mapped COs. Note that AIS 7 corresponds to the example in B. E) Histogram of CO density along AIS depth. Almost all COs are in the initial 30 *µ*m. F) Minimum distance from ChC synapses to COs within the same AIS and where ChC synapses are computationally mapped to the same depth and orientation on each other AIS. Solid lines are median, dashed lines are interquartile intervals. G) Minimum distance from ChC synapses to COs from the same AIS, but where ChC synapses are rotated in orientation. The solid line is the mean value across all AIS, the shaded line is the 95% confidence interval from bootstrapping. Stars indicate significance of a t-test measuring difference from zero rotation. In F and G, comparison is by t-test. *: p<0.05, **: p<0.01, ***: P<0.001.

## Discussion

In this study we analyzed the chandelier input to pyramidal cells from the fine anatomy of the subcellular organization of their synapses to the structure of their network architecture and their functional role *in vivo.* Both structural and functional results suggest that chandelier cells act collectively to deliver a common inhibitory signal to their targets. This global inhibitory signal covaries with arousal and it is distributed differentially to each individual pyramidal cell with connectivity rules governed by at least three independent factors. These results indicate that this cell-type specific connection is likely regulated by a multitude of rules that we are just beginning to understand.

### Structured variability and implications for function

In agreement with previous observations in mouse somatosensory cortex (Inan et al., 2013), we find that ChCs formed synapses with nearly all PyCs in L2/3. However, the degree to which a PyC received input from chandelier axons exhibited strong postsynaptic heterogeneity. This wide distribution was already noted in early studies from cat and monkey (DeFelipe et al., 1985; Fairén and Valverde, 1980), but has remained largely unexamined. We found that three basic structural properties relating only to a PyC perisomatic region could explain approximately half the variance in the number of synaptic connections. These properties fell into two categories: those that affected the density of nearby ChC axons, and those that affected the connection likelihood to those nearby axons. How might each of these properties reflect the functional effect of ChC on cortical circuits?

The density of ChC axons within L2/3 decreased with depth, and this was reflected in a reduced ChC synapse count onto deeper PyCs. A similar result was also found in mouse prelimbic cortex using electrophysiology from ChCs and PyCs labeled by long-range projection target (Lu et al., 2017). In that work, shallow amygdala-projecting PyCs had a higher connection probability with nearby ChCs than deeper cortical-projecting PyCs. In mouse V1, graded differences in the projections of L2/3 PyCs to higher order visual areas have also been observed (Kim et al., 2018). Depth-dependent variability in ChC inhibition could thus be well-posed to allow differential state-dependent inhibition of distinct long-range excitatory subnetworks.

Increased ChC input was associated with two structural factors: larger PyCs soma size and increased total somatic synapse count. The effect of size is consistent with a compensatory effect. If a particular target inhibitory postsynaptic potential is desired by the PyC, more ChC synapses would be necessary to balance the lower input resistance of a larger cell. Such a homeostatic target point for ChC inhibition has been suggested by experiments that manipulated PyC activity and observed compensatory changes in the number of ChC boutons (Pan-Vazquez et al., 2020) as well as the location and size of the AIS (Wefelmeyer et al., 2015). The relationship between somatic and AIS input also suggests a cell-specific target point for inhibition, potentially to appropriately match excitation on a cell-by-cell level (Vogels and Abbott, 2009; Vogels et al., 2011). Since somatic input is principally GABAergic and ChCs do not target the soma, this suggests that different inhibitory cell types respond to the same (or correlated) inhibitory target points. As different interneurons are active at different times, this would be an effective way to ensure that the appropriate amount of inhibition is available across conditions and states. Interestingly, both factors relating to target point manifest as increased connection probability with nearby ChC axons, raising the possibility that both structural factors engage similar mechanisms to form or prune connections.

One limitation of the dataset is that all cells and neuronal processes were truncated by the boundaries of the imaged volume, although this volume is among the largest cortical datasets of its kind currently available. We took care to account for this truncation throughout our analysis by only studying neurons and compartments which we could map completely. However, this could introduce some biases or faulty assumptions. First, we have undersampled the deeper part of L23, since those axons left the volume closer to the soma. PyCs in deep L3 could have different properties that we did not measure here. It is also possible that the decrease in ChC input with depth is more of a categorical step than a linear decline, but this would require additional data in L3. Second, we assumed that every ChC axon targeting the same AIS came from a different ChC. Light-level morphology suggests it is rare for multiple axonal branches of a ChC to target the same AIS, but it is possible. This would not affect the overall synapse count analysis, but it could overestimate the number of distinct connections. However, given the strong correlation between the number of distinct connections and overall ChC synapse count, the interpretation of the data would be largely unchanged. Future work in a larger dataset could help not only resolve these issues but allow the use of richer data about both whole-cell morphology and a more complete synaptic network. Moreover, given that simple perisomatic features already accounted for approximately half the variance in ChC synapse count, we suspect that such information would reveal additional factors related to ChC input, for example functional activity, PyC subtypes, or excitatory network structure.

### Functional role of ChCs

The functional role of ChCs has been far less studied than their structure and it is not known what conditions drive ChC activity. Whether in neocortex or in the hippocampus, Chandelier cells have been shown to fire in a brain-state dependent manner in anesthetized animals (Klausberger et al., 2003; Massi et al., 2012), however their function in awake animals has not been described before. Here, we used functional imaging of ChCs in awake behaving animals and observed strong, synchronous bouts of activity during periods of pupil dilation, a key sign of an active arousal state (McGinley et al., 2015; Reimer et al., 2016; Vinck et al., 2015). Interestingly, during pupil dilation ChCs are tonically active, as are layer 1 cholinergic axon projections (Reimer et al., 2016), which are known to innervate ChC dendrites (Lu et al., 2017). Activity in visual cortex changes significantly during arousal, with PyCs generally reducing their spontaneous activity and increasing their signal to noise ratio (Vinck et al., 2015). A subclass of VIP interneurons that are active during locomotion have been strongly implicated in this modulation (Fu et al. 2015). Our data suggest that ChC also contribute to the high arousal state. Assuming ChCs have an inhibitory role under these conditions, this suggests that ChCs add a significant extra source of inhibition to some — but not all — PyCs during the high arousal state, in contrast to the combination of disinhibition (Fu et al., 2014; Karnani et al., 2016) and direct inhibition (Kuljis et al., 2018) from VIP neurons. L2/3 PyCs in visual cortex show a broad diversity of activation and suppression of activity during dilations of the pupil (Stringer et al., 2016; Vinck et al., 2015). The heterogeneity of ChC inhibition strength could be one circuit mechanism underlying this diversity if, for example, adjusting the amount of ChC input a PyC receives would modulate the degree to which it is sensitive to arousal-driven depolarizations.

### A role for global inhibition in recurrent cortical circuits

As we show here, ChCs are active during arousal states, which are known to be associated with an increase in the signal to noise ratio in pyramidal cells (Vinck et al. 2015). One hypothesis lies on the connectivity motif that we extracted from the anatomy, where the recurrently connected excitatory network receives a common signal from a pool of inhibitory ChCs. In addition, ChCs also receive input from the same local excitatory network, a connectivity arrangement that resembles the soft winner-take-all (sWTA) motif originally proposed by (Amari and Arbib, 1977) and others (Douglas et al., 1995; Hahnloser et al., 2000; Maass, 2000). A basic feature of the sWTA motif is its ability to amplify the response of the subset of neurons that receive the strongest input, while the responses of the others are suppressed due to the shared common inhibition (which can, in addition, be dynamically regulated by either the local excitatory network or additional external inputs). When on top of an input signal one adds a common excitatory signal (such as arousal) to all the neurons of the network, this common signal acts as a gain modulator that selectively increases the overall signal to noise (Douglas and Martin, 2007).

### Subcellular targeting at the AIS

The result that ER cisternae are colocalized with synapses on a lateralized portion of the AIS, raises the possibility that ChC inputs are located on the AIS in order to biochemically separate them from other perisomatic synapses. This biochemical separation could facilitate the implementation of alternative plasticity rules (Pan-Vazquez et al., 2020; Schlüter et al., 2017), through different calcium dynamics influenced by the ER (reviewed in Berridge, 1998), or through differential sorting of molecular components into the AIS (reviewed in Leterrier, 2018). The functional significance of the lateralization of the ER cisternae and synaptic input remains unclear, but could relate to distinct mechanisms that affect neuronal excitability via calcium mediated modulation of ion channels. There is much evidence that different ion channels have precise regulation in terms of location along the AIS (reviewed in (Leterrier, 2018)), but there is little exploration of their distribution around its circumference.

### A framework to map cell type connectivity rules

We are only beginning to understand the logic of connectivity between cortical cell types. The underlying rules of connectivity between neurons depend not only on pre- and postsynaptic cell types, but geometry, morphology, and function, as well as, as we have shown here, various intrinsic properties of the cells involved. Indeed, it is likely that connections between different cell types will depend in different ways on different factors. Large-scale cortical connectomics (Bock et al., 2011; Kasthuri et al., 2015; Lee et al., 2016; Motta et al., 2019), particularly in the context of physiological measurements (Bock et al., 2011; Lee et al., 2016), offers the promise to examine the connections between identified cell types in the context of other structural, functional, and network measurements. The use of electron microscopy is crucial, as it is the only way to densely map synaptic connectivity with simultaneous measurements of the detailed morphology and individual anatomical features for any cell. The connection from ChCs to L23 PyCs is a model case for investigating the nuances of cell-type specific connectivity, as the strong anatomical specificity makes a complete map easier to acquire. We anticipate that this approach, applied to the upcoming generation of millimeter-scale EM volumes, will be a powerful tool to uncover the wiring rules across the complete diversity of cortical cell types.

## Acknowledgments

We thank Wenjing Yin for re-imaging of sections with the EM. We thank Rob Young for managing the stitching and alignment pipeline at the Allen Institute for Brain Science (AIBS). We thank John Philips, Sill Coulter and the Program Management team at the AIBS for their guidance for project strategy and operations. We thank Hongkui Zeng, Ed Lein, Christof Koch and Allan Jones for their support and leadership. We thank the Manufacturing and Processing Engineering team at the AIBS for their help in implementing the EM imaging and sectioning pipeline. We thank Brian Youngstrom, Stuart Kendrick and the Allen Institute IT team for support with infrastructure, data management and data transfer. We thank the Facilities, Finance, and Legal teams at the AIBS for their support on the MICrONS contract. We thank the Neurosurgery and Behavior and the Transgenic Colony Management teams at the AIBS for the preparation of mice for calcium imaging of Chandelier cells. We thank Stephan Saalfeld for help with the parameters for 2D stitching and rough alignment of the dataset. We would like to thank the “Connectomics at Google” team for developing Neuroglancer and computational resource donations. We also would like to thank Amazon and Intel for their assistance. We thank S. Koolman, M. Moore, S. Morejohn, B. Silverman, K. Willie, and R. Willie for their image analyses, Garrett McGrath for computer system administration, and May Husseini and Larry and Janet Jackel for project administration. We thank Ueli Rutishauser, Ahmed El Hady and G. Ocker for advice and feedback. Supported by the Intelligence Advanced Research Projects Activity (IARPA) via Department of Interior/ Interior Business Center (DoI/IBC) contract numbers D16PC00003, D16PC00004, and D16PC0005. The U.S. Government is authorized to reproduce and distribute reprints for Governmental purposes notwithstanding any copyright annotation thereon. Disclaimer: The views and conclusions contained herein are those of the authors and should not be interpreted as necessarily representing the official policies or endorsements, either expressed or implied, of IARPA, DoI/IBC, or the U.S. Government. HSS also acknowledges support from NIH/NINDS U19 NS104648, ARO W911NF-12-1-0594, NIH/NEI R01 EY027036, NIH/NIMH U01 MH114824, NIH/NINDS R01NS104926, NIH/NIMH RF1MH117815, and the Mathers Foundation. We thank the Allen Institute for Brain Science founder, Paul G. Allen, for his vision, encouragement and support.

## Author Contributions

Conceptualization A.A.B, C.S.M, F.C, N.M.C and R.C.R; Software (Alignment) G.M., F.C;, R.T., T.M.,W.M.S and W.W.; Software (Analysis infrastructure) C.S.J., C.S.M., F.C and S.D..; Software (Data interaction and viewing) M.C., N.K. and W.M.S.; Software (Cloud data storage) W.M.S.; Software (Proofreading System) J.Zu., N.K. and S.D.; Formal Analysis A.N., B.H., C.S.M., F.C., J.Zh. and S.D.; Investigation (Calcium Imaging from Chandelier Cells) J.Zh.; Investigation (Calcium Imaging from EM volume;) E.F, and J.R..; Investigation (Histological preparation for EM), J.B., M.T. and N.M.C.; Investigation (Sectioning sample for EM), A.A.B and A.L.B.; Investigation (EM Dataset Generation), A.A.B, A.L.B, D.J.B., J.B. and N.M.C.; Investigation (Alignment) D.I., G.M., I.T., R.T, T.M., Y. L. and W.W.; Investigation (Neuron Reconstruction) A.Z., I.T., J.Zu., J.W., K.L., M.C., N.K., R.L., S.P., and T.M.; Investigation (Synapse Detection) J.W and N.L.T.; Investigation (Sectioning sample for EM) A.A.B., and A.L.B; Investigation (Biophysical Modelling) A.N. and C.A.; Data Curation (Proofreading) A.A.B., A.L.B., C.S.M., F.C., N.M.C. and S.D., Writing-original C. A., C.S.M, F.C., H.S.S., N.M.C and R.C.R. Writing – Review & Editing A.A.B, A. N., C. A., C.S.M, F.C., H.S.S., J.Zh., N. L. T., N.M.C and R.C.R; Visualization C.S.M. and F.C., Supervision A.S.T., C.A., H.S.S., N.M.C. and R.C.R., Project administration A.S.T., L.B., S.S., H.S.S., N.M.C. and R.C.R., Funding Acquisition A.S.T., H.S.S., N.M.C. and R.C.R.

## Supplemental Figures

**Supplemental Figure 1, related to Figure 1 and Figure 3.**
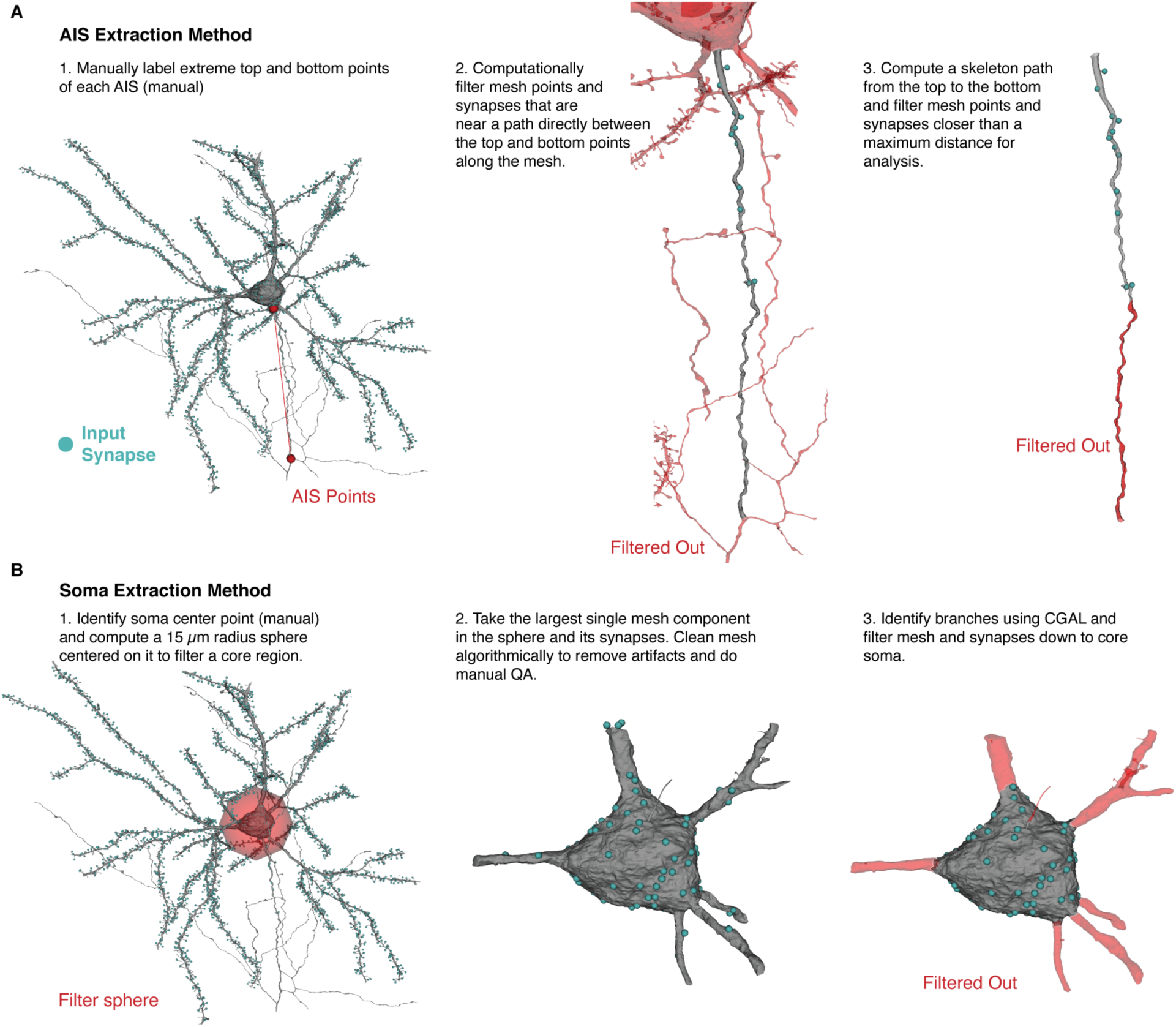
Computational steps involved in defining the AIS and soma compartments and their synapses. A, AIS extraction. From left to right: (1) We start with a full soma mesh (gray) with all automatically detected inputs (cyan) and a pair of manually annotated AIS points (red). (2) We find the initial axonal mesh region that is near a direct path the two points along the surface of the mesh and keep that mesh region and the synapses on it (Mesh region: grey, synapses: cyan dots), while filtering out the rest of the mesh and its synapses (red). (3) We specify the initial region of the AIS up to a maximum distance by computing a simple “skeleton”, i.e. the shortest path along the mesh from the top-most (i.e. shallowest) point of initial axonal mesh to the bottom-most (i.e. deepest) point. We measure the distance along the skeleton from the top. For every point of the axonal mesh, we find the closest point on the skeleton path and remove those mesh points (and their associated synapses) whose closest skeleton point is beyond the maximum distance (red). B, Soma Extraction. From left to right: (1) Starting from the same whole mesh, we manually identified the approximate centroid of each PyC soma and defined a 15 *µ*m radius sphere around it (red), which is sufficient to capture the soma of all neurons. (2) We filtered out the mesh outside of the sphere and algorithmically omitted artifacts and components from axonal or dendritic branches that come into the sphere from outside. The remaining soma mesh and its synapses (gray, cyan dots respectively) includes proximal neurites. (3) The computational vision package CGAL identifies regions with substantial changes in radius. We used this to define the mesh region at the soma and keep the associated synapses.

**Supplemental Figure 2, related to Figure 2.**
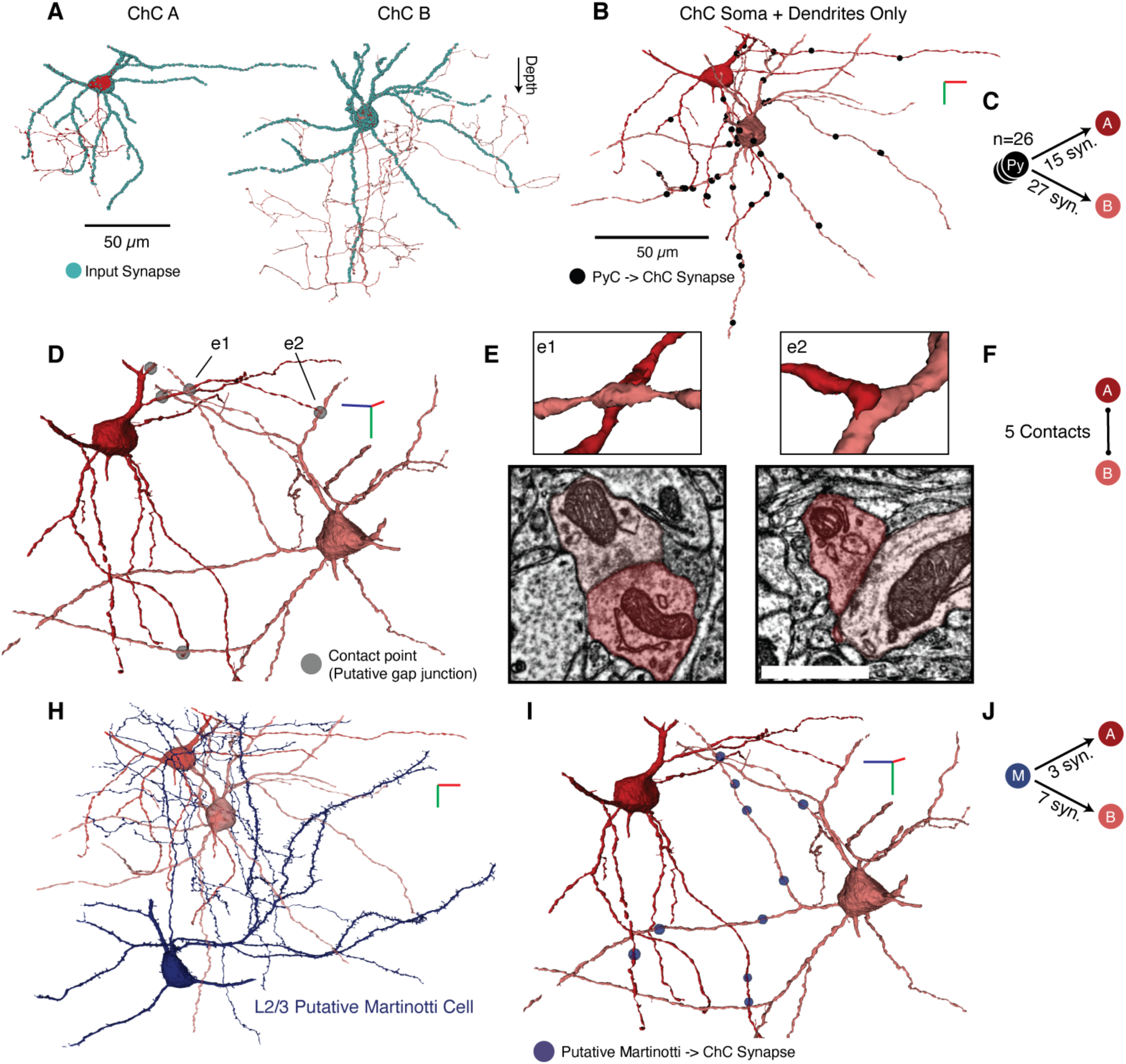
Dendritic input into ChCs. A, We found two ChCs (denoted A and B, respectively) with cell bodies and substantial dendrites in the volume, shown here individually for clarity. Axons and dendrites are both incomplete, as they leave the imaged volume boundaries. Both dendrites and soma are covered with synaptic input (cyan dots). Note that the synapse-free region on the soma of cell A is exactly where it is directly apposed to another cell body. B, Both ChCs receive synaptic input from the pyramidal cells (black dots) in the volume across their dendrites. Individual PyCs placed between 1-4 synapses on target ChCs, although that is a lower bound. C, In total, 26 PyCs made 42 synapses with the two ChCs, suggesting widespread feedback from the local excitatory network. D, Putative gap junctions between ChCs. We found five contact points (gray circles) with electron-dense membrane structures between ChC dendrites, four at terminal points. E, Detailed 3d contact morphology (top) and EM imagery of the two contacts. Scale bar is 500 nm. Note the dense clouds at the contact sites. F, Graph representation of the putative gap junction contacts. H, A putative L2/3 Martinotti cell (based on morphology) makes synapses across both ChC dendrites. J, Graph representation of connections onto each ChC from the Martinotti cell (M). In panels B, D, H, and I the red/green/blue 3d axes are shown to specify rotation angle (green is towards white matter). Each line of the axis is 10 *µ*m.

**Supplemental Figure 3, related to Figure 3.**
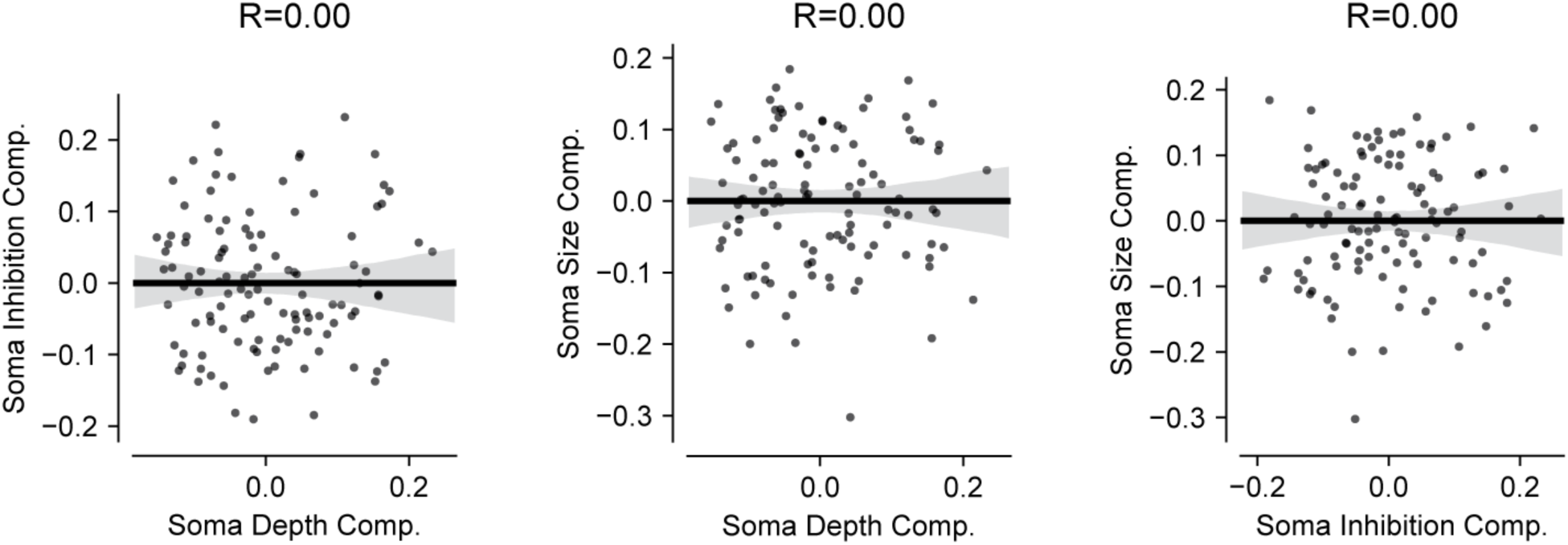
ICA finds uncorrelated structural components for the PyCs. Each dot represents one PyC. Lines indicate least squares linear fit, shaded region is a 95% confidence interval estimated from bootstrap resampling.

**Supplemental Figure 4, related to Figure 3.**
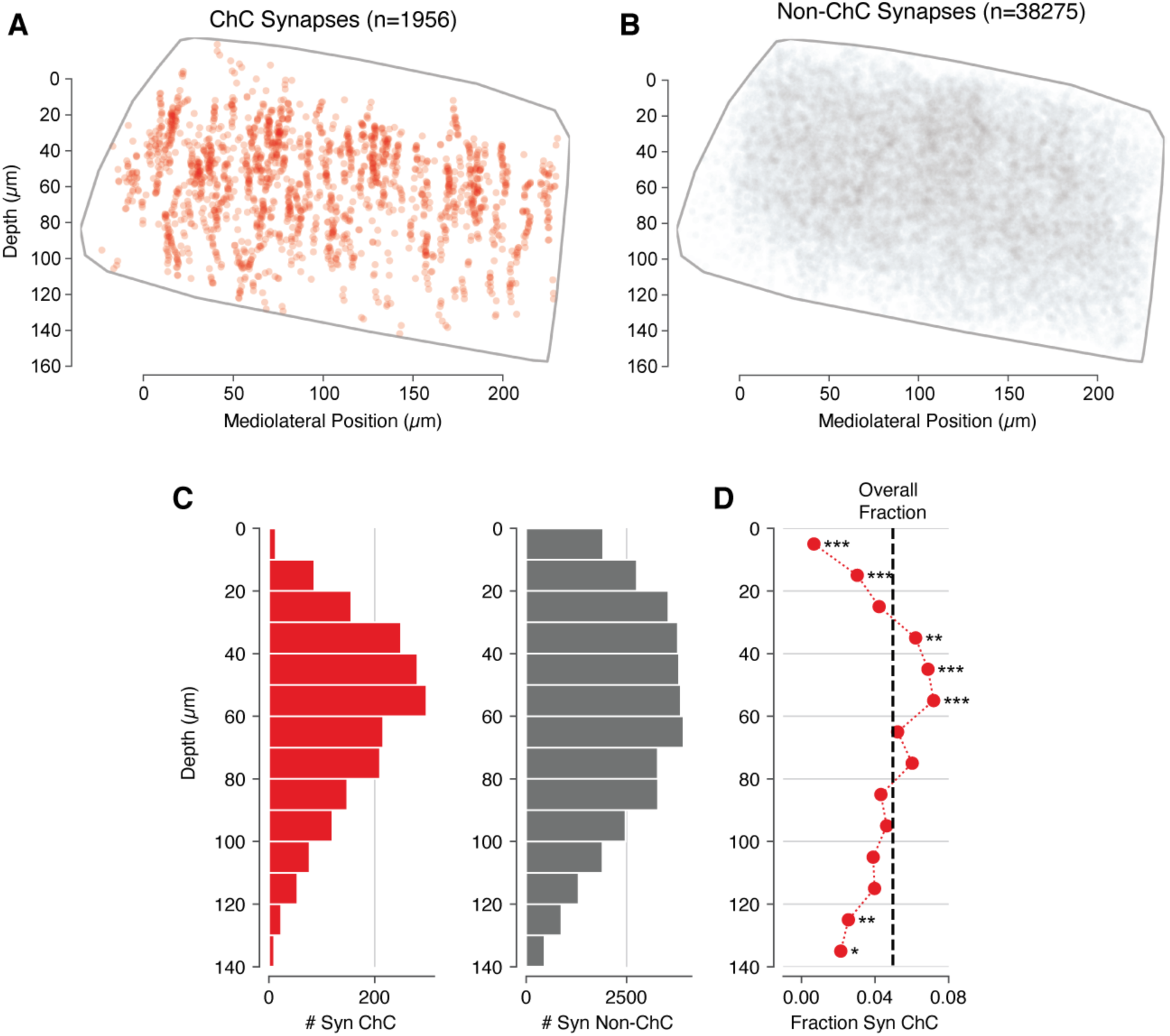
Distribution of ChC synapses in L2/3. A) All output synapses from measured ChCs (red), whether onto measured AISes or not), within the volume. Gray outline indicates the approximate segmentation boundary, pia is up. B, Distribution of all outputs synapses from observed non-ChC cells (grey), onto any target, as in A. We use this as an estimate of where synapses could reasonably fall in the dataset conditional on having any synapses on one of the AISs that we considered. C, Density of synapses as a function of depth for the ChCs (left) and non-ChCs (right). D, Red line, fraction of observed synapses in a given depth bein above that are from ChC (i.e. # ChC Syn. / (# ChC Syn. + # non-ChC Syn.)). Black line: Same fraction, measured across all depths. Stars indicate a significant difference from the overall average (*: p<0.05, **: p<0.01, ***: p<0.001, Fisher exact test). ChC synapses are overrepresented in a band between 30–60 *µ*m from the top of the dataset, approximately at the L1/L2 border).

**Supplemental Figure 5, related to Figure 4.**
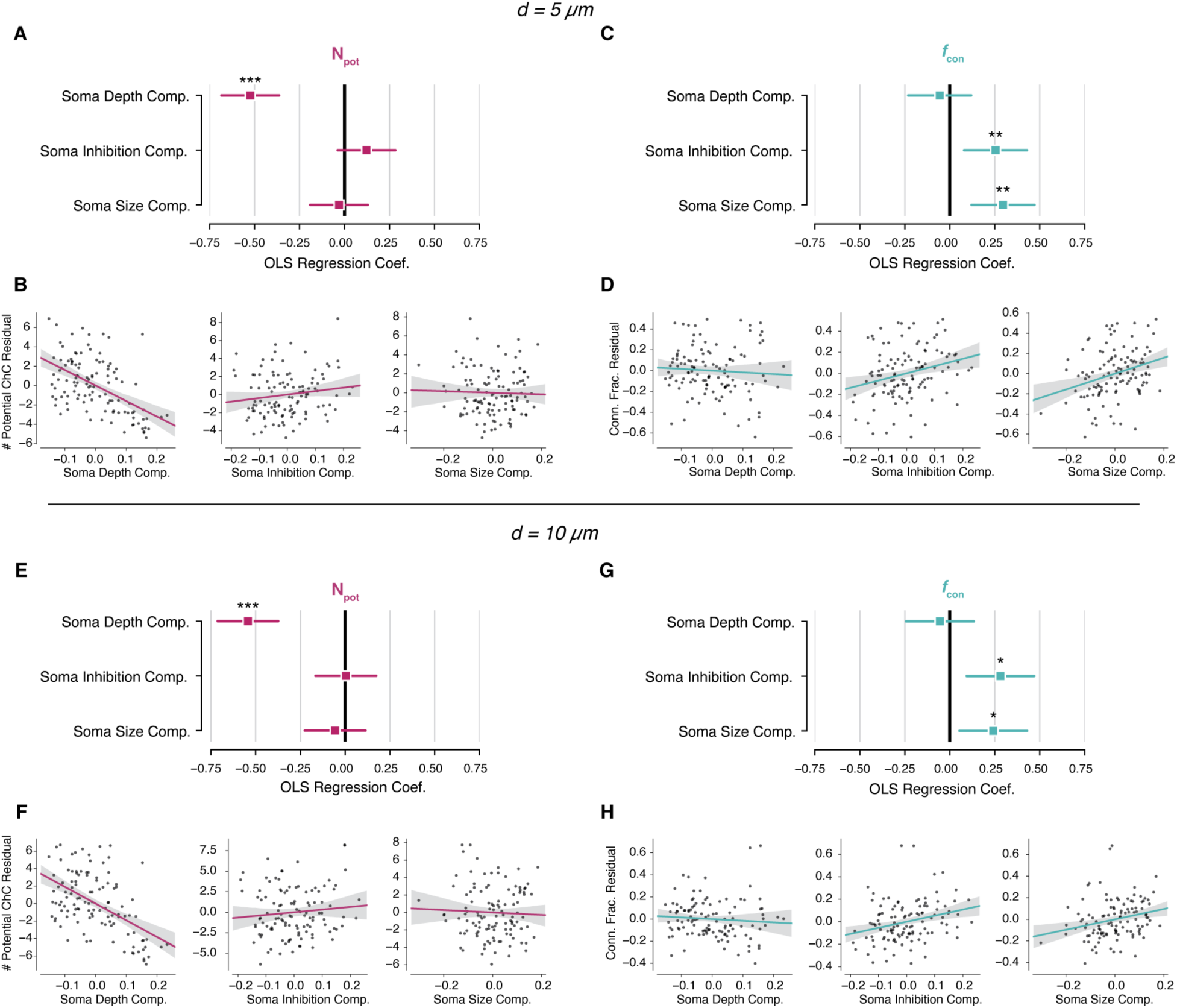
OLS analysis of potential connectivity for alternative distance thresholds. For d=5 *µ*m: A), OLS coefficients for number of potential connections, as in Figure 4H. B), Each component plotted against the residuals for potential connections after fitting the other two components, as in Figure 4I. Black line indicates least squares fit, gray shade indicates a 95% confidence interval based on bootstrapping. C), OLS coefficients for connectivity fraction, as in Figure 4J. D), Each component plotted against the residuals for connectivity fraction after fitting the other two components, as in Figure 4K. E–H), same as A–D but for a potential connection distance threshold of 10 *µ*m.

**Supplemental Figure 6, related to Figure 5.**
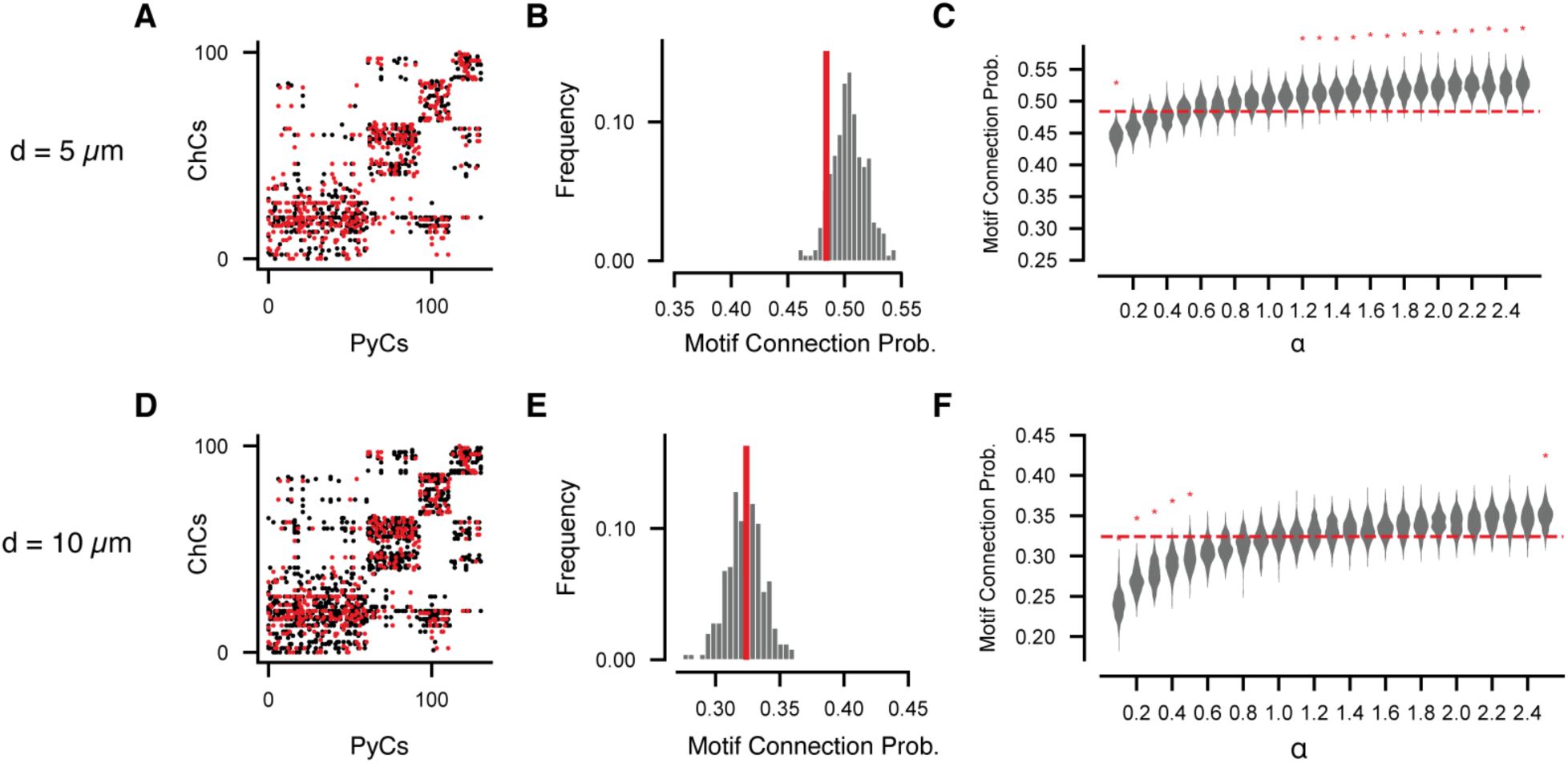
Robustness of potential motif analysis for different potential distance thresholds. Using a distance threshold of 5 *µ*m: A), Bipartite adjacency matrix of potential (black) and actual (red) connections, using the same node ordering as Figure 5A. B), Observed potential bifan motif probability (red line) compared to shuffled graphs (equivalent to Figure 5C). C), Comparison of observed motif connection probability (red line) with shuffled graphs with different weightings (equivalent to Figure 5E). For a distance threshold of 10 *µ*m: D), E), and F) follow as in A, B, and C respectively.

**Supplemental Video 1, related to Figure 1**. A rendering of the EM reconstructions from this dataset demonstrating the mapping of ChC inputs onto L2/3 PyCs. Video begins with four gray PyCs with only their somatic regions and AIS region shown. An individual pink ChC fragment is slowly revealed over time, as the reconstruction is followed along to all the locations that it synapses onto. Note, portions of that axon which are far from the four PyCs are excluded from the rendering for clarity. Then a second, purple axon fragment is revealed in the same fashion. Third, all the ChC fragments which synapses onto these four PyCs are revealed simultaneously, each with their unique color. Finally, the scene pans out to reveal all the PyCs in this structural dataset, and all of the ChCs reconstructed in the dataset are revealed in red.

## Methods

### Animal preparation for electron microscopy

All animal procedures were approved by the Institutional Animal Care and Use Committee at the Allen Institute for Brain Science.

Neurophysiology data acquisition was conducted at Baylor College of Medicine. To aid in registration of optical physiology data to EM data, a wide field image of the cranial window visualizing the surface vasculature was provided in addition to a volumetric image stack of the vasculature, encompassing the region of tissue where the neurophysiology dataset was acquired. The vasculature was imaged by injecting a red fluorescent dye into the bloodstream of the mouse, allowing blood vessels and cell bodies to be imaged simultaneously by 2-photon microscopy. Mice were then transferred to the Allen Institute in Seattle, where they were kept in a quarantine facility for 1 to 3 days after which they were euthanized and perfused.

### Perfusion

After induction of anesthesia with isoflurane, the appropriate plane of anesthesia was checked by a lack of toe pinch reflex and the animals were transcardially perfused with 15 ml 0.15 M cacodylate buffer (EMS, Hatfield, PA, pH 7.4) followed by 30 ml fixative mixture containing 0.08 M cacodylate (pH 7.4), 2.5% paraformaldehyde (EMS), 1.25% glutaraldehyde (EMS) and 2 mM calcium chloride (Sigma). The perfusion solution was based on the work of (Hua et al., 2015). Once the brain was removed it was placed into the same fixative solution to post-fix for 16 to 72 hours at 4 °C.

### Identifying neurophysiological region for further electron microscopy processing

To accurately identify and isolate the region of cortical tissue where the neurophysiology dataset was imaged, we labeled and imaged the surface and descending vasculature. After perfusion of the animals and excision of the brain, the surface of the cortex was imaged using differential contrast lighting to visualize the surface vasculature of visual cortex where the cranial window had previously been. We manually marked fiduciary points around this region of cortex to aid identification of the previously imaged cortical site after vibratome sectioning. The brain was washed in CB (0.1 M cacodylate buffer pH 7.4) and embedded in 2% agarose. The agarose was trimmed and mounted for coronal sectioning in a Leica VT1000S vibratome; successive 200 μm thick slices were taken until the entire region of cortical tissue previously demarcated by manual markings was sectioned. During this procedure, we also acquired blockface images of each brain slice.

After vibratome sectioning, the wide field image of the cranial window and the vasculature stack were co-registered with the images of the surface vasculature from the brain surface using TrakEM2 (Cardona et al., 2012). From the blockface images, we could readily identify the fiduciary marks made on the brain surface. A volumetric representation of the cortical surface of the blockface images and the fiduciary marks was constructed in TrakEM2 and the orientation and position of the vibratome sections were aligned to the surface vasculature images by affine registration of the fiduciary points. This allowed for determination of the vibratome slices that contained the previously imaged region of cortical tissue.

To map the neurophysiology imaged site within the coronal slice, we next mounted the slices under coverglass in CB and acquired 10x images of the entire hemisphere of the slice, and 20x image stacks covering the cortical tissue surrounding the potential imaged site using a Zeiss AX10 ImagerM2 upright light microscope. These 10x and 20x stacks were co-registered in trakEM2, and these images allowed us to visualize the descending vasculature within the coronal sections. We next generated a volumetric rendering of the vasculature stack provided by Baylor College of Medicine using microView (Parallax Innovations). From this rendering, we could reslice and visualize the descending vasculature from the previously imaged site. Corresponding vasculature landmarks were identified between the *in-vivo* imaged site and the light microscopy coronal slice stacks. These landmarks were used to map the extent of the previously imaged tissue site to the coronal slice. The coronal sections containing the imaged site were then selected for histological processing (see below).

### Electron microscopy Histology

The histology protocol used here is based on the work of (Hua et al., 2015), with modifications to accommodate different tissue block sizes and to improve tissue contrast for transmission electron microscopy (TEM).

Following several washes in CB (0.1 M cacodylate buffer pH 7.4), the vibratome slices were treated with a heavy metal staining protocol. Initial osmium fixation with 2% osmium tetroxide in CB for 90 minutes at room temperature was followed by immersion in 2.5% potassium ferricyanide in CB for 90 minutes at room temperature. After 2 × 30 minute washes with deionized (DI) water, the tissue was treated with freshly made and filtered 1% aqueous thiocarbohydrazide at 40 °C for 10 minutes. The samples were washed 2 × 30 minutes with DI water and treated again with 2% osmium tetroxide in water for 30 minutes at room temperature. Double washes in DI water for 30 min each were followed by immersion in 1% aqueous uranyl acetate overnight at 4°. The next morning, the samples in the same solution were placed in a heat block to raise the temperature to 50° for 2 hours. The samples were washed twice in DI water for 30 minutes each, then incubated in Walton’s lead aspartate pH 5.0 for 2 hours at 50 °C in the heat block. After double washes in DI water for 30 minutes each, the slices were dehydrated in an ascending ethanol series (50%, 70%, 90%, 3 × 100%) 10 minutes each and two transition fluid steps of 100 % acetonitrile for 20 minutes each. Infiltration with acetonitrile:resin dilutions at 2p:1p (24 h), 1p:1p (48 h) and 1p:2p (24 h) were performed on a gyratory shaker. Samples were placed in 100% resin for 24 hours, followed by embedment in Hard Plus resin (EMS, Hatfield, PA). The samples were cured in a 60 °C oven for 96 hours.

In order to evaluate the quality of samples during protocol development and before preparation for large scale sectioning, the following procedure was used for tissue mounting, sectioning and imaging. We evaluated each sample for membrane integrity, overall contrast and quality of ultrastructure. For general tissue evaluation, adjacent slices and tissue sections from the opposite hemisphere, processed in the same manner as the ROI slice, were cross-sectioned and thin sections were taken for evaluation of staining throughout the block neighboring the region of interest.

### Ultrathin Sectioning

The tissue block was trimmed to contain the neurophysiology recording site which is the region of interest (ROI) then sectioned to 40 nm ultrathin sections. For both trimming and sectioning a Leica EM UC7 ultramicrotome was equipped with a diamond trimming tool and an Ultra 35 diamond knife (Diatome USA) respectively. Sectioning speed was set to 0.3 mm/sec. Eight to ten serial thin sections were cut to form a ribbon, after which the microtome thickness setting was changed to 0 nm in order to release the ribbon from the knife edge. Then, using an eyelash superglued to a handle, ribbons were organized to pairs and picked up as pairs to copper grids (Pelco, SynapTek, 1.5 mm slot hole) covered by 50nm thick LUXFilm support (Luxel Corp., Friday Harbor, WA).

### Electron microscopy imaging

The imaging platform used for high throughput serial section imaging is a JEOL-1200EXII 120kV transmission electron microscope that has been modified with an extended column, a custom scintillator, and a large format sCMOS camera outfitted with a low distortion lens. The column extension and scintillator facilitate an estimated 10-fold magnification of the nominal field of view with negligible impact on resolution. Subsequent imaging of the scintillator with a high-resolution, large-format camera allows the capture of fields-of-view as large as 13×13um at 4nm resolution. As with any magnification process, the electron density at the phosphor drops off as the column is extended. To mitigate the impact of reduced electron density on image quality (shot noise), a high-sensitivity sCMOS camera was selected and the scintillator composition tuned in order to generate high quality EM images within exposure times of 90 - 200 ms (Yin et al., 2019).

### Proofreading and Annotation of Volumetric Imagery Data

We used a combination of Neuroglancer (Maitin-Shepard, https://github.com/google/neuroglancer) and custom tools to annotate and store labeled spatial points. In brief, we used Neuroglancer to simultaneously visualize the imagery and segmentation of the 3d EM data. A custom branch of Neuroglancer was developed that could interface with a “dynamic” segmentation database, allowing users to correct errors (i.e. either merging or splitting neurons) in a centralized database from a web browser. Neuroglancer has some annotation functionality, allowing users to place simple annotations during a session, but does not offer a way to store them in a central location for analysis. We thus built a custom cloud-based database system to store arbitrary annotation data centered associated with spatial points that could be propagated dynamically across proofreading events. Annotations were programmatically added to the database using a custom python client and, in relevant cases, after parsing temporary Neuroglancer session states using custom python scripts. These spatial points and their associated data (e.g. synapse type, cell body ID number, or cell types) were linked to stored snapshots of the proofreading for querying and reproducible data analysis. All data analyzed here came from the “v183” snapshot.

### Visualization and Analysis of Mesh Data

Neuronal meshes were computed by Igneous (https://github.com/seung-lab/igneous) and kept up to date across proofreading. Meshes were analyzed in a custom python library, MeshParty (https://github.com/sdorkenw/MeshParty), that extends Trimesh (https://trimsh.org) with domain-specific features and VTK (https://www.vtk.org) integration for visualization. In cases where skeletons were used, we computed them with a custom modification of the TEASAR algorithm (Sato et al., 2000) on the vertex adjacency graph of the mesh object implemented as part of MeshParty. In order to associate annotations such as synapses or AIS boundary points with a mesh, we mapped point annotations to the closest mesh vertex after removing artifacts from the meshing process.

### AIS Identification and Extraction

To get a handle on the contents of the EM volume, we manually identified every cell body in the dataset manually (N=458) and assigned a unique point at the approximate center of the soma for each. We then manually assigned a coarse cell type (excitatory, inhibitory, or glia) to each based on morphology. To identify the AIS of excitatory neurons, we manually placed points at the top and bottom of excitatory neurons that were thought to have a largely completely AIS in the volume. To extract the mesh vertices associated with the AIS, we computed the path distance between each mesh vertex and the top and bottom points (*d*_*top*_ and *d*_*bottom*_ respectively), the distance between top and bottom points (*d*_*AIS*_), and kept those vertices that satisfied 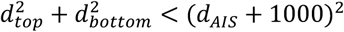. Distances were measured in nm using Dijkstra’s algorithm as implemented in SciPy (https://www.scipy.org). The constant padding term allows the mesh definition to wrap around the AIS smoothly.

To extract the initial 37 microns of the AIS, we skeletonized each AIS (see previous section) and computed which mesh vertices were closest to each skeleton vertex. We filtered out all mesh vertices associated with the skeleton vertices more than 37 microns of the top skeleton tip. The threshold was chosen to balance keeping as many distinct PyCs as possible while covering as much of the ChC input domain as possible.

### Chandelier Cell Type Classification and Proofreading

Cell typing of AIS inputs was performed across several rounds of proofreading and annotation. Every neurite presynaptic to an AIS was evaluated starting from its synapses. The main step was to evaluate other synapses from the same axon. The compartment (AIS, dendrite, or soma) was trivial to determine via manual inspection from the automated segmentation, even without labels or proofreading. If any of those synapses targeted dendrites or soma at subsequent points along the axon, the axon was labeled non-ChC at the seed synapses. Attaching the annotation to this point would allow any potential distant splitting of the axon due to proofreading to remain unlabeled. In contrast, axons that exhibited multi-bouton cartridges characteristic of ChCs and only targeted AISs were labeled as ChCs and proofread completely, extending tips and splitting segmentation errors. Because of the large number and size of these cells, comprehensive proofreading of non-ChC was beyond the scope of the project. However, to account of the possibility of ChC axons that had been erroneously merged into non-ChC axons by the automated segmentation, we evaluated every non-ChC synapse after completing an initial round of axon classification. Non-ChC axons forming multi-synaptic contacts onto any AIS were also given additional scrutiny. A small number of axonal fragments that targeted AIS near the edge of the volume had few synapses overall and were more difficult to classify. In those cases, we used bouton morphology, tight clustering with established ChC boutons, and in some instances following the axonal process in imagery outside the segmented region to cartridges.

### Soma and AIS Mesh Structural Properties

The automated meshing process introduced artifacts that made measuring spatial properties like surface area require special processing. For example, if part of the nucleus was segmented separately, this would introduce extra mesh faces to the neuron. In order to extract a clean surface for cell somata, we extracted a cutout of the voxel segmentation within 15 microns of the approximate cell body center that contained both the soma and initial part of proximal neurites. Boundary expansion and contraction for the segmentation was performed to fill small gaps caused by image and segmentation artifacts and the mesh was recomputed using the marching cubes algorithm. We then used CGAL (https://www.cgal.org) surface mesh segmentation to identify proximal neurites and leave only the core soma mesh for measurements. Manual quality control of resulting meshes was done to ensure that the analysis only included cell bodies that were completely contained in the volume and were free of remaining artifacts. Surface area was computed by summing the area of mesh faces. Synapse count was computed from those synapses associated with the core soma mesh.

We computed AIS radius from the mesh via ray tracing. From each point on the AIS skeleton, a ray was sent toward the opposite side and the intersection with the opposite point was used to determine the diameter at that point. The AIS can emerge from either the soma or a dendrite, which could influence the radius of the most proximal part of the AIS due to the transition between compartments. To avoid being affected by transition, we ignored the initial 5 *µ*m and averaged the radius based on skeleton vertices between 5-38 *µ*m along the AIS.

The imagery dataset was sectioned so that the y-axis was approximately aligned with the pia-to-white-matter dimension, but exact alignment was not possible at the data collection stage. To more accurately measure depth within the data, we rotated the coordinate system to align with the average vector from AIS top to AIS bottom across all cells.

### Structural Components Analysis

We could compute complete AIS and soma features for 113 cells in the data. Most cells that were excluded had soma that touched the edge of the volume and thus were only partially reconstructed. Pearson correlations between structural features were found using Scipy with a Holm-Sidak multiple test correction. To address the underlying correlations, we used the FastICA implementation in scikit-learn to do a components analysis of the structural properties. We selected three components, as the first three components of a principal components analysis (an approximation and bound on ICA explained variance) account for 84% of the variance and three ICA components were clearly interpretable. FastICA is a stochastic algorithm, so we ran it many times and selected the most robust solution. Components were multiplied by −1 if needed to make the largest element positive.

To look at the relationship between structural components and ChC input, we used ordinary least squares regression on z-scored counts of synaptic input, number of connections, and average synapses per connection. Coefficients and confidence intervals were computed with Statsmodels (citation) and p-values were adjusted with a Holm-Sidak multiple test correction.

### Potential Synapse Analysis

We defined a potential connection as a ChC axon whose mesh came within a given distance threshold of a truncated AIS mesh (as described above). Distances were measured with the Scipy implementation of the k-d tree data structure. We tested distance thresholds between 5–15 *µ*m and saw qualitatively similar results for 5-10 *µ*m. Since the number of true connections was irrespective of distance, connectivity threshold was strictly non-increasing with increased distance. For a distance threshold of 15 *µ*m, connectivity fraction showed a negative trend with depth, suggesting that the influence of depth on potential connections became a factor at that length scale due to density of nearby axons, and we report only the results from the lower range of thresholds. For some AISs near the edge of the volume, the region around them might reach beyond the edge of the segmentation, resulting in a potential undercounting of potential connections. To avoid including these AIS in our analysis, we computed the fraction of voxels within the distance threshold that were within the segmented data. If more than 10% of the voxels were outside of the segmentation for a given distance threshold, the cell was omitted from that analysis.

### Network Motif Analysis

Based on the definition of potential connection above, we generated two bipartite networks from ChCs onto PyCs, one based on the potential connectivity and one based on actual synaptic connectivity. By definition, the actual connectivity network is a strict subset of the potential network. To investigate the clumpiness of ChC targeting, we looked at the bifan motif in the bipartite network from ChC to PyCs. A bifan is defined as the motif comprising two source nodes (ChCs) and two target nodes (PyCs) and four edges, here such that two ChC target the same two PyCs. We generalized the concept to include the concept of potential synapse, defining a “potential bifan” as a bifan where one edge was a potential connection and the other three were actual connections. For every potential connection, we evaluated if it was part of a potential bifan and then if it was an actual bifan. The bifan connectivity fraction was then the ratio of these two numbers.

We next developed a method to randomize the actual network within the potential network. Starting from the observed actual network, we iterated through each actual connection in a random order and at each step we: 1) removed the actual connection from the graph, leaving it only a potential connection and 2) for the target AIS, picked a new potential, but not already connected, ChC (including the just-disconnected one). Each step preserves the degree distribution of the AIS, but not the ChC, which we chose to reflect the apparent postsynaptic influence on ChC input and relative uniformity of ChC axons. We iterate through the network five times, shuffling each edge each time. To add a bias to the shuffling, at the step of picking a new actual connection we evaluated which potential connections could complete a bifan motif. For each ChC connection *i* it was given a weight *w*_*i*_ *= α* if it was a potential bifan and *w*_*i*_ *=* 1 otherwise. A connection was then selected with probability proportional to its weight. The result is that for *α* > 1, connectivity will be clumpier with bifans being generated more often than chance, while for *α* < 1, connectivity will be more dispersed, with bifans being generated less often than chance.

### Cisternal Organelle Annotation and Analysis

To annotate the COs, we selected ten PyCs from intentionally from across the distribution of total ChC synapse count. An expert neuroanatomist worked through every 3-4 sections of the data, placing annotation points on ER with stacked cisternae. Cisternae points were placed within the extent of the organelle every several sections, creating a point cloud throughout.

To compute the location of the COs, we mapped the stacked cisternae points to the closest mesh index on the AIS. This left both clear clumps associated with COs and a few diffuse outliers due to ambiguous ultrastructure and the mapping procedure. To filter the data down to just the clumps, we used a density-based clustering algorithm DBSCAN (citation) as implemented in scikit-learn, where the distance between points was computed along the mesh vertices and edges.

While synapse detection associated a location with every synapse, we found that there was a bias in this location away from *en face* parts of the synapse active zone and onto the transversely cut, resulting in an axis-aligned bias of AIS synapse locations. To more accurately assign a location to each synapse, we associated each synapse with the contact site between the presynaptic ChC mesh and the postsynaptic AIS mesh. The contact site was computed by finding nodes of the AIS mesh within 150 nm of the ChC mesh and clustering with DBSCAN (Ester et al., 1996) using precomputed distances along the mesh surface, which generated puncta-like clusters for each bouton contact. Each synapse could then be associated with a given puncta, and the center of the puncta was computed by identifying the node with the highest average distance along the mesh surface from other nodes within the puncta.

For each synapse, we computed its depth and orientation of the mesh vertex associated with it. Depth was measured as the distance along the skeleton from the top to the skeleton node closest to the synapse mesh vertex. Orientation was measured by computing a slice of the AIS mesh centered at the synapse and spanning 400 nm along the pia-to-white matter direction (the y-axis of the dataset). The AIS mesh points in this slice were projected into 2d and their convex hull was used to estimate the local outline of the AIS. The angle of the vector from the center of the convex hull to the synapse was used for the orientation, with an orientation angle of 0 corresponding to the positive x-axis direction. To map vertices across AISes, we used their depth and orientation values of each synapse to compute the vertex on the target AIS with the most similar orientation that was also within 200 nm of the same depth.

### Generating biophysically detailed all-active models

The Allen Cell-Types database (http://celltypes.brain-map.org) contains 1920 *in vitro* whole cell patch clamp recordings and 485 biocytin filled digital reconstructions from neurons in mouse primary visual cortex for a variety of transgenic lines. The all-active single neuron models are constrained by these two data modalities from the same cell, namely, electrophysiology – voltage responses under standardized set of protocols and morphology – diameter and length of each segment within the tree. We distribute voltage gated Sodium (Na^+^), Potassium (K^+^) and Calcium (Ca^++^) conductances across the entire morphology, specifically, we use the following channels Ih, NaT, NaP, KT, KP, Kv2, Kv3.1, SK, Im, CaLVA, CaHVA, with the assumption that these ion channels are expressed uniformly along each major morphological sections: soma, axon, apical and basal dendrites. These parameters and the passive membrane properties (membrane capacitance cm, axial resistance Ra, leak conductance g_pas, reversal potential e_pas) construct the multicompartmental model of the neuron, thereby forming the variable vector (n = 43) for the optimization. To avoid inconsistencies in the axonal reconstructions such as isolated segments (or absence of axon altogether), we replace the axon with a 60 *µ*m long, 1 *µ*m diameter initial segment. We use a multiobjective optimization framework (Druckmann et al., 2007) where features such as action potential (AP) amplitude, width, spike frequency, steady state voltage etc., are extracted from individual traces and the deviation (z-score) between experiment and model feature at a specific stimulus becomes one of the objectives. For our purposes we have used python toolbox BluePyOpt (Van Geit et al., 2016) that offer evolutionary algorithms (Fortin et al., 2012) to solving multiobjective optimization problems, with NEURON 7.5 (Hines and Carnevale, 1997) under the hood to simulate each model spawned out of the parameter explorations. To get a handle on the computation we have designed a 3 stage workflow where we progressively introduce new channels to the circuit, i.e., first only fit the passive parameters with features from subthreshold experimental traces, next add hyperpolarization activated channel Ih with sag (Hogan and Poroli, 2008) related features to the objective, and finally equip the circuit at its full complexity by introducing the rest of the channels and fit the conductance densities with both spiking and subthreshold trace features. Throughout the workflow the Indicator based evolutionary algorithm (IBEA) adds 512 new offsprings (new models) to the population at each generation with Stage 0,1 evolved up to 50 generations with 1 seed and Stage 2 continued for 200 generations with 4 independent seeds. On a 256 × 2.2 GHz Intel Xeon E5-2630v4 distributed cluster with 150 Gb of maximum process memory the optimization of a single cell takes 26 ± 11 hours. Overall we have used 15 distinct features across the three stages extracted with the eFEL library (Van Geit et al., 2018). Our models capture axonal AP initiation, an important aspect of biophysical realism. To add this constraint, we append the boolean feature checkAISInitiation, part of the eFEL library, at the final stage of the optimization. This involves calculating the AP onset at axon and soma and adding a heavy penalty to the models for which somatic AP precedes axonal AP. This also requires allowing a less restrictive maximal density for the transient Na+ current and lower action potential threshold at the axon compared to the soma. At the conclusion of the 3 stages, the workflow outputs 40 models sorted according to the sum of all objectives, for each experiment. We use the ‘best model’, i.e., the model with least training error for downstream analysis of synaptic integration properties.

### Simulating single neuron models under synapses

For simulating the single neuron models under synaptic inputs, we have used Brain Modeling Toolkit (bmtk) (Gratiy et al., 2018) with NEURON 7.5 simulation environment. In this study we have used all-active models for 4 cells with ids: 477127614, 571306690, 584254833, 382982932 from the Allen Cell-Types database. For Figure 7E,F we have simulated each model for 1 sec with 50 excitatory synapses over 25 connections (2 synapses per connection) with Poisson spike train rates ranging from 0 to 500 Hz. The number of inhibitory synapses on the axon initial segment (AIS) of the modeled PyC and their distribution is adopted from EM data and the frequency of the incoming inhibitory spike train is held constant at 100 Hz. For each cell and number of inhibitory synapses to evaluate the resultant ‘fi curve’ (e) we run 8 different repetitions at each excitation rate. We aggregate the spike count output for these simulations and group them according to the amount of inhibition and perform one sided pairwise Wilcoxon signed-rank test at 5% false discovery rate.

For the simulations in Figure 7G both excitatory and inhibitory spike trains are sampled from a gaussian rate function with the maximal rate of 500 Hz and 200 Hz respectively. The peak of the excitation is varied at 5 ms intervals with 10 repetitions to capture the variability in response and the total simulation window is 1 sec. The number of excitatory synapses and their distribution (25*2) remains unchanged. For the pairwise comparison, we once again use one-sided Wilcoxon signed-rank test at 5% FDR. The free parameters in these simulations, such as the maximal excitation, inhibition rate, or number of excitatory synapses, are selected such that the cells represented by the biophysical models operate at a similar activation regime.

### Surgery and animal preparation for in vivo imaging

In total, three Vipr2-IRES2-Cre mice (1 male, 2 female) were used in this study. The surgery included a stereotaxic viral injection and a cranial window/head-plate implantation. During the injection, a glass pipette back-loaded with AAV virus was slowly lowered into the superficial layer of left V1 (3.8 mm posterior 2.7 mm lateral from bregma, 0.3 mm below pia) through a burr hole. 5 minutes after reaching the targeted location, 50 or 100 nL of virus was injected into the brain over 10 minutes by a hydraulic nanoliter injection system (Nanoject III, Drummond). The pipette then stayed for an additional 10 minutes before it was slowly retracted out of the brain. AAV1 (or AAV5)-CAG-FLEX-GCaMP6f (addgene: 100835-AAV1 or AAV5, titer 2×10^13^ vg/ml) were used for functional imaging and AAV9-CAG-FLEX-eGFP (addgene: 51502-AAV9, titer 2.28×10^13^ vg/ml, 1:10 dilution) was used for structure imaging. Immediately after injection, a titanium head-plate and a 5 mm glass cranial window were implanted over left V1 following the Allen Institute standard procedure protocol (de Vries et al., 2020; for detailed protocol see Goldey et al., 2014) allowing *in vivo* two-photon imaging during head fixation.

After surgery, the animals were allowed to recover for at least 5 days before retinotopic imaging with intrinsic signal during anesthesia (for detailed retinotopic protocol, see (Juavinett et al., 2017)). After retinotopic mapping, animals were handled and habituated to the imaging rig for two additional weeks before *in vivo* two-photon imaging (de Vries et al., 2020).

All experiments and procedures were approved by the Allen Institute Animal Care and Use Committee.

### Histology for viral expression

To characterize the Cre expression pattern, Vipr2-IRES2-Cre mice injected by Cre-dependent eGFP or GCaMP viruses were perfused and brains collected. Briefly, mice were anesthetized with 5% isoflurane and 10 ml of saline (0.9% NaCl) followed by 50 ml of freshly prepared 4% paraformaldehyde (PFA) was pumped intracardially at a flow rate of 9 ml/min. Brains were immediately dissected and post-fixed in 4% PFA at room temperature for 3-6 hours and then overnight at 4 °C. After fixation, brains were incubated in 10% and then 30% sucrose in PBS for 12-24 hours at 4 °C before being cut into 50 *µ*m sections by a freezing-sliding microtome (Leica SM 2101R). Sections from V1 were mounted on gelatin-coated slides and cover-slipped with Prolong Diamond Antifade Mounting Media (P36965, ThermoFisher). For GCaMP labeled tissue, sections were processed with antibody staining before mounting. During antibody staining, sections containing LGN and V1 were blocked with 5% normal donkey serum and 0.2% Triton X-100 in PBS for one hour, incubated in an anti-GFP primary antibody (1:5000 diluted in the blocking solution, Abcam, Ab13970) for 48-72 hours at 4 °C, washed the following day in 0.2% Triton X-100 in PBS and incubated in a Alexa-488 conjugated secondary antibody (1:500, 703-545-155, Jackson ImmunoResearch) and DAPI.

The sections were then imaged with Zeiss AxioImager M2 widefield microscope with a 10x/0.3 NA objective. Fluorescence from antibody enhanced GCaMP and mRuby3 were extracted from filter sets Semrock GFP-1828A (excitation 482/18 nm, emission 520/28 nm, dichroic cutoff 495 nm) and Zeiss # 20 (excitation 546/12 nm, emission 608/32 nm, dichroic cutoff 560 nm), respectively. A subset of sections was imaged with 2-photon microscope (Scientifica 2PIMS, objective: Nikon 16XLWD-PF, 16x/0.8 NA, laser: Coherent Chameleon Ultra II, excitation wavelength: 920nm, emission filter: 470-558 nm band-pass) to obtain 3d axon morphology with high resolution (pixel size: 0.47 *µ*m).

### *In vivo* two-photon imaging

When recovery, retinotopic mapping, habituation was finished (usually more than three weeks after initial surgery), V1 cells labeled with GCaMP6f were evident in superficial layers (50 – 200 *µ*m) through the cranial window. Calcium activities from those cells were imaged by two-photon excitation using a custom microscope and 940 nm illumination by a Ti:sapphire laser (a Spectra-Physic Insight X3), focused with a 16×/0.8 NA objective (Nikon N16XLWD-PF). This scope has the ability to correct optical aberration (adaptive optics) and quickly switch focal depth by modulating the beam wavefront with a liquid crystal spatial light modulator (SLM, Meadowlark Optics, HSP-512, Liu et al., 2018). With this scope, we recorded calcium activities from planes at 3 different depths (16 or 32 *µ*m apart in depth) in single imaging sessions. The plane at each depth was sequentially imaged at about 37 Hz and the volume rate was about 12 Hz and aberrations out of the objective were corrected (with 512 x 512 pixels resolution at LSM). Emitted light was first split by a 735 nm dichroic mirror (FF735-DiO1, Semrock). The short-wavelength light was filtered by a 750 nm short-pass filter (FESH0750, Thorlabs) and a 470-588 nm bandpass emission filter (FF01-514/44-25, Semrock) before collected as GCaMP signal. Image acquisition was controlled using Vidrio ScanImage software (Pologruto et al., 2003). To maintain constant immersion of the objective, we used gel immersion (Genteal Gel, Alcon).

All imaging sessions were performed during head fixation with the standard Allen Institute Brain Observatory *in vivo* imaging stage (de Vries et al., 2020).

### 2-photon Image preprocessing

The recorded 2-photon movies for each imaging plane were motion-corrected using rigid body transform based on phase correlation by a custom-written python package (https://github.com/zhuangjun1981/stia/tree/master/stia, (Zhuang et al., 2017)). To generate regions of interest (ROIs) for boutons, the motion-corrected movies were further downsampled by a factor of 3 and then processed with constrained non-negative matrix factorization (CNMF) (Pnevmatikakis et al., 2016), implemented in the CaImAn python library (Giovannucci et al., 2019). These ROIs were further filtered by their size (ROIs smaller than 23.4 *µ*m^2^ or larger than 467.5 *µ*m^2^ were excluded), position (ROIs within the motion artifacts were excluded). Since the labeled cells distributed sparsely, there were no overlapping pixels among ROIs. For each retained ROI, a neuropil ROI was created as the region between two contours by dilating the ROI’s outer border by 1 and 8 pixels excluding the pixels within the union of all ROIs. The calcium trace for each ROI was calculated by the mean of pixel-wise product between the ROI and each frame of the movie, and its neuropil trace was calculated in the same way using its neuropil ROI. To remove the neuropil contamination, the neuropil contribution of each ROI’s calcium trace was estimated by a linear model and optimized by gradient descendent regression with a smoothness regularization (Zhuang et al., 2017; de Vries et al., 2020).

### Pupil area and locomotion speed extraction

During each imaging session, the locomotion speed and a movie of the animal’s right eye were simultaneously recorded following the Allen Brain Observatory standard protocol (de Vries et al., 2020, Zhuang et al., 2017). To extract pupil area, the contour of pupil in each frame was extracted with adaptive thresholding (Zhuang et al., 2017) by custom-written python package “eyetracker” (https://github.com/zhuangjun1981/eyetracker).

For each imaging session, a comprehensive collection of data including metadata, visual stimuli, all preprocessing results, final calcium traces, locomotion speed, and pupil area was generated in Neurodata Without Borders (nwb) 1.0 format with “ainwb” package (https://github.com/AllenInstitute/nwb-api). The analysis pipeline (imaging preprocessing, pupil/locomotion analysis, and nwb packaging) were performed by a custom-written python package “corticalmapping” (https://github.com/zhuangjun1981/corticalmapping).

### Correlation Analysis

We computed pairwise correlations between the activities of all recorded cells for each imaging session during spontaneous periods. For the chandelier cell recordings, we analyzed cells across the three different imaging planes. For the non-chandelier cell types (VIP, SST, and PV), we used data recorded from a single imagine plane in visual area VISp. This data was downloaded through the publicly available Allen Brain Observatory using the AllenSDK (0.16.3). We first downsampled the corrected fluorescence traces from 30 Hz to 12 Hz (using the scipy resample function) to match the sampling rate of the chandelier cell recordings. We show the distribution of correlation coefficients across all simultaneously recorded cell pairs separated by cell type, and report summary statistics comparing the mean correlation coefficients by cell type. We compute statistical significance between the chandelier and non-chandelier cell correlation values using the non-parametric Mann-Whitney U test.

To compute correlations between cell activity and other behavioral covariates, for each cell, we computed the correlation between its calcium activity and pupil area / locomotion speed. We compute statistical significance between the mean correlation values for running and pupil diameter using the Wilcoxon signed-rank test.

## Resources

Segmented and annotated EM data will be available online upon submission, as will all analysis scripts.

